# Mapping gene regulatory networks of primary CD4^+^ T cells using single-cell genomics and genome engineering

**DOI:** 10.1101/678060

**Authors:** Rachel E. Gate, Min Cheol Kim, Andrew Lu, David Lee, Eric Shifrut, Meena Subramaniam, Alexander Marson, Chun J. Ye

**Author notes:** These authors contributed equally to this work (Rachel E. Gate, Min Cheol Kim). Corresponding author. (C.J.Y.).

## Abstract

Gene regulatory programs controlling the activation and polarization of CD4^+^ T cells are incompletely mapped and the interindividual variability in these programs remain unknown. We sequenced the transcriptomes of ~160k CD4^+^ T cells from 9 donors following pooled CRISPR perturbation targeting 140 regulators. We identified 134 regulators that affect T cell functionalization, including *IRF2* as a positive regulator of Th_2_ polarization. Leveraging correlation patterns between cells, we mapped 194 pairs of interacting regulators, including known (e.g. *BATF* and *JUN*) and novel interactions (e.g. *ETS1* and *STAT6*). Finally, we identified 80 natural genetic variants with effects on gene expression, 48 of which are modified by a perturbation. In CD4^+^ T cells, CRISPR perturbations can influence *in vitro* polarization and modify the effects of *trans* and *cis* regulatory elements on gene expression.

---

CD4^+^ T cells display an incredible degree of functional diversity during adaptive immune responses classically characterized by the expression of canonical cytokines that create systemic inflammatory responses (e.g. Th_17_) (*1*–*4*), signal and recruit B cells (e.g. Th_1_ and Th_2_) (*5*–*11*), and induce tolerance in the tissue microenvironment (e.g. T_reg_) (*12*, *13*). Recently, the ability of CD4^+^ T cells to directly kill infected and tumor cells have also received renewed attention (*14*–*16*). While the functionalization of CD4^+^ T cells is predominantly determined by the polarization of naive T cells (T_naive_), recent results have suggested a high degree of variation within and plasticity between canonical subtypes (*17*). Indeed, we (*18*, *19*) and others (*20*–*22*) have shown that human circulating CD4^+^ T cells are composed of a mixture of canonical and non-canonical populations with significant interindividual variability in both the proportion and gene expression of each population (*18*).

Key regulators of CD4^+^ T cell differentiation and polarization have been mapped in mice utilizing pooled and arrayed genetic screens. For example, pooled knockdown screens with RNA interference have identified *Ppp2r2d* as a key regulator of T cell proliferation (*23*) and mapped two self-reinforcing, mutually antagonistic modules of regulators that drive Th_17_ differentiation (*24*). More recently, a pooled genome wide CRISPR screen paired with bulk RNA sequencing identified *Trappc12*, *Mpv17l2*, and *Pou6f1* as regulators of both Th_2_ activation and differentiation (*25*). Despite these insights, mapping gene regulatory programs that underlie human CD4^+^ T cell state transitions is incomplete and the intra- and inter-individual variability in these programs remain largely unknown.

Recent advances in the integration of droplet-based single cell RNA-sequencing (dscRNA-seq) and CRISPR/Cas9-mediated genome engineering has created new opportunities to assess the functional consequences of genetic perturbations in primary human T cells at an unprecedented molecular and cellular resolution (*26*–*28*). Here, we integrate single guide RNA (sgRNA) lentiviral infection with Cas9 protein electroporation (SLICE) and multiplexed dscRNA-seq (mux-seq) to screen the effects of 140 regulators in primary CD4^+^ T cells across nine donors. By linking each sgRNA with the transcriptomes of hundreds of heterogeneous CD4^+^ cells, we map regulators that affect the activation and polarization of specific T cell subsets. By further leveraging the coexpression patterns across single cells, we define novel gene regulatory relationships between pairs of regulators and their downstream targets. Finally, by incorporating donor genetics, we identify instances where genetic effect on gene expression is modified by CRISPR perturbations. Our work demonstrates that systematic analyses using multiplexed single-cell genomics and genome engineering is a powerful approach to map the gene regulatory networks that govern the functionalization of primary human T cells and characterize the intra- and inter-individual variability in these networks.

## Results

### CRISPR perturbation screen in activated CD4^+^ T cells across donors

To map gene regulatory programs that underlie the polarization and activation of human CD4^+^ T cells, we performed sgRNA lentiviral infection with Cas9 protein electroporation (SLICE) (*28*) followed by multiplexed single-cell RNA-sequencing (mux-seq). Primary CD4^+^ T cells were isolated from peripheral blood mononuclear cells (PBMCs) and activated *in vitro* as previously described (*29*, *30*) (Fig. 1A; **Methods**). Activated cells were transfected with 280 sgRNAs targeting 140 regulators that were either highly expressed (top quartile from bulk RNA-seq) or have binding sites that were differentially accessible (from bulk ATAC-seq) in activated CD4^+^ T cells (*18*) (Fig. 1B; **Table 1**; **Methods**). Following Cas9 electroporation and multiple rounds of selection and proliferation, activated CD4^+^ T cells (*31*) were pooled across 9 donors and profiled using the 10X Chromium platform in 16 wells resulting in 320,708 cell-containing droplets and 16,750 reads/droplet (**Fig. S1**-**3**; **Methods**). To maximize the probability of detecting sgRNAs in cells, we further amplified and sequenced the sgRNA transcripts from the resulting 10X cDNA library to near saturation as previously described (*32*) (98% compared to 63% in the 3’ tagged library; Fig. 1C; **Methods**).

**Figure 1:**
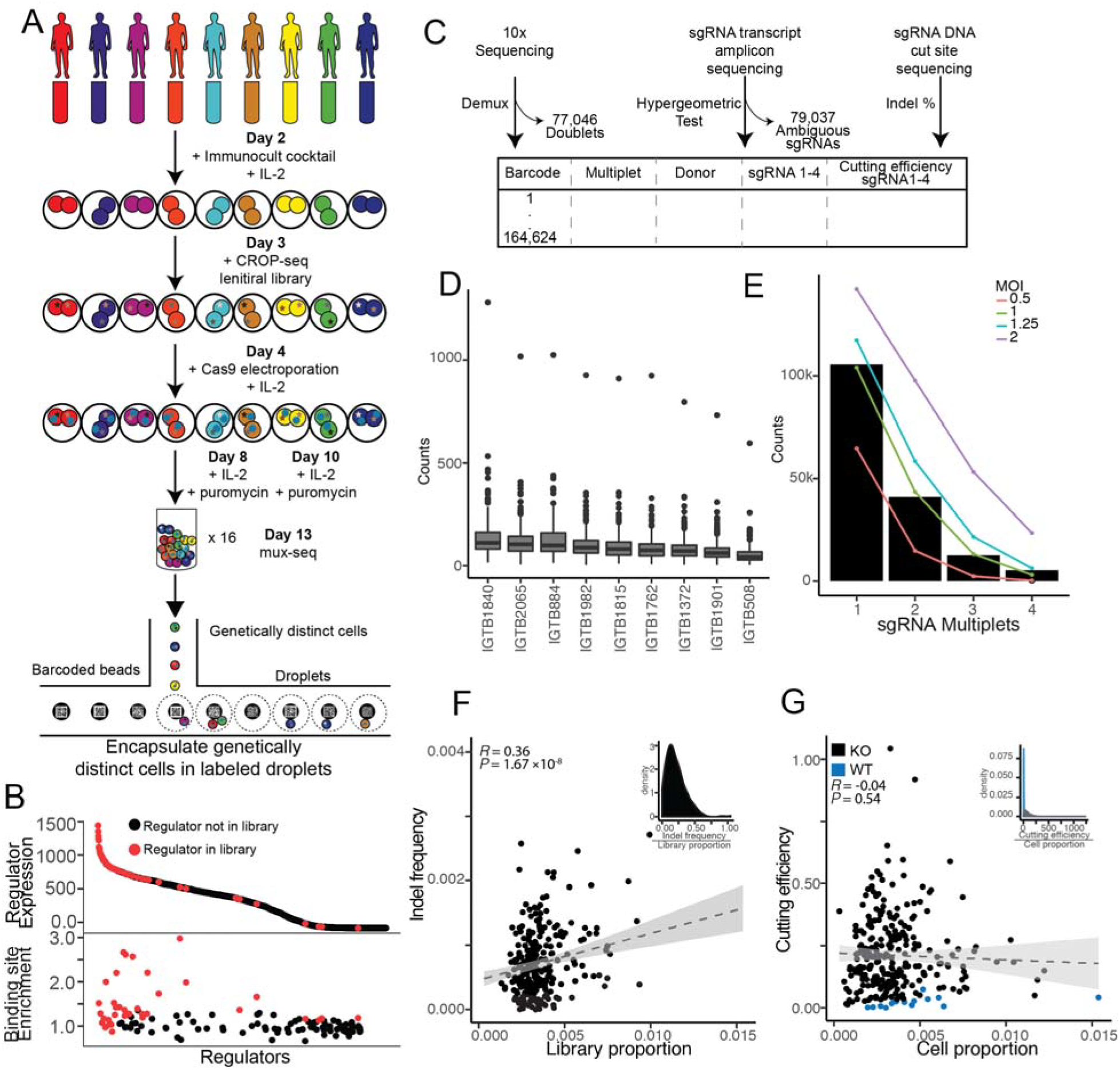
CRISPR perturbation screen in activated CD4^+^ T cells across donors. **(A)** Experiment overview. **(B)** Unbiased identification of candidate regulators from transcript abundance (top) and accessibility of binding sites (bottom) in activated CD4^+^ T cells (*18*). Targeted regulators are in red and all other regulators in the human genome are in black. **(C)** Data processing overview of 10X single-cell RNA-sequencing, sgRNA amplicon sequencing, and target loci DNA sequencing. **(D)** Total number of cells expressing each sgRNA per donor. **(E)** Observed distribution of cells with 1-4 sgRNA (black bars) and expected Poisson distributions at an MOI of 0.5 (pink), 1 (green), 1.25 (blue), and 2 (purple). **(F)** For each sgRNA, cutting efficiency (inset) is estimated as the ratio of indel frequency at the targeted locus (y-axis) and sgRNA frequency in the pool (x-axis).(**G)** For each wildtype (WT: blue) and knockout (KO: black) sgRNA, ratio (inset) of cutting efficiency (y-axis) and the proportion of cells expressing the sgRNA (x-axis).

After filtering for doublets using exonic SNPs (77,046 - 24%) and cells with ambiguous sgRNA assignments (79,037 cells - 25%; **Methods**), 164,623 cells were kept for subsequent analyses corresponding to 18,291±4,571 cells per donor and 882±522 cells per sgRNA (Fig. 1C, D; **Table 2**; **Methods**). Of these, 105,664, 40,960, 12,675, and 5,324 cells contained one, two, three, or four sgRNAs, resulting in an estimated multiplicity of infection (MOI) of 1 (Fig. 1E, **S4**).

To assess the cutting efficiency of each sgRNA, we sequenced the sgRNA pool and DNA of the edited cells from each donor by targeted amplification of 268/280 loci (**Methods**). The insertion and deletion (indel) frequencies at each targeted locus and coverage of the corresponding sgRNA in the pool are expectedly correlated (*R* = 0.36, *P* < 1.68×10^−8^, Fig. 1F) and the average ratio between these two quantities - defined as the cutting efficiency - is 21%±15% (Fig. 1F **inset**). We defined 14 sgRNAs as uncut negative controls (WT) where the ratios between the cutting efficiencies and proportion of cells containing each sgRNA are 1.645 standard deviations below the mean (z-score < −1.645, *P* < 0.05) (Fig. 1G, **S5**; **Methods**). In all, the integration of SLICE and mux-seq is an efficient and cost-effective strategy for pooled screening and profiling of primary human T cells across many donors.

### Heterogeneity of activated CD4^+^ T cells

Because CD4^+^ T cells dynamically migrate to and egress from tissues through circulation, cells isolated from PBMCs likely represent a functionally diverse population of cells reflective of the specific immunological state of an individual. This is supported by previous functional genomic analyses of activated primary CD4^+^ cells demonstrating marked heterogeneity within and variability between individuals in the chromatin and expression profiles that overlap signatures from multiple sorted populations including Th_1_s, Th_2_s, and Th_17_s (*18*, *22*, *33*).

To assess the heterogeneity of activated CD4^+^ T cells using dscRNA-seq, we clustered 164,623 cells into 10 Leiden clusters (*34*) with each cluster containing on average 16,462 cells (maximum: 65,720; minimum: 127; Fig. 2A, B). We identified 2,189 differentially expressed genes in at least one cluster (624±691 per cluster) and annotated each cluster based on the most highly expressed markers. We found a *CD27*^+^/*CCR7*^+^ naive T cell population (T_naive_) (*35*, *36*) (65,720 cells, 40%), and three effector populations including *IL5*^+^/*IL17RB*^+^/*GATA3*^+^ Th_2_s (*37*–*45*) (1,969, 1.1%), *IFNG*^+^ Th_1_s (*42*, *46*, *47*) (8,022 cells, 4.8%), and *PRF1*^+^/*GNLY*^+^/*NKG7*^+^ cytotoxic cells (T_cyto_) (*16*) (37,960, 23%) (Fig. 2C, D). We also identified three activated populations (Th_stim_) defined by the expression of *HMGB2* and *STMN1 (48, 49)* and distinguished from each other by the expression of histone markers (Th_stim,histone_, 19,920 cells, 12%), cell cycling markers *PTTG1 (50, 51)* and *KIAA0101 (52–54)* (Th_stim,cycling_, 16,905 cells, 10%), and naive markers *CCNB1 (55)* (T_stim,naive_, 12,345 cells, 7.4%). We also found a proliferating population (T_prolif_) that expressed genes associated with tumor progression, including *FXYD5 (56–58)*, *LIMD2 (59, 60)*, and *PFDN5 (61)* (903 cells, 0.05%). Finally, we identified two small clusters likely to be transitional, as they are intermediates in lineage trajectory (*62*) space either between naive and cytotoxic cells (T_naive→cyto_ 752 cells, 0.04%) or between naive and stimulated cells (T_naive→stim_: 172 cells, 0.01%; **Fig. S6**). To validate these annotations, for each cluster, we correlated the average log fold change in expression of upregulated genes (with respect to all other clusters) to the bulk RNA-seq expression profiles across 45 reference circulating immune populations (*22*) (**Methods**). This approach assigned 8/10 clusters to their expected reference population (Fig. 2E, **Methods**). The frequency of each cluster was generally consistent across donors (average pairwise *R* = 0.94±0.04; Fig. 2F, **S7**). These results demonstrate that multiplexed single-cell RNA-sequencing of activated CD4^+^ T cells recapitulate the expected T cell subpopulations obtained from sorted PBMCs and enables estimates of donor variability in T cell composition.

**Figure 2:**
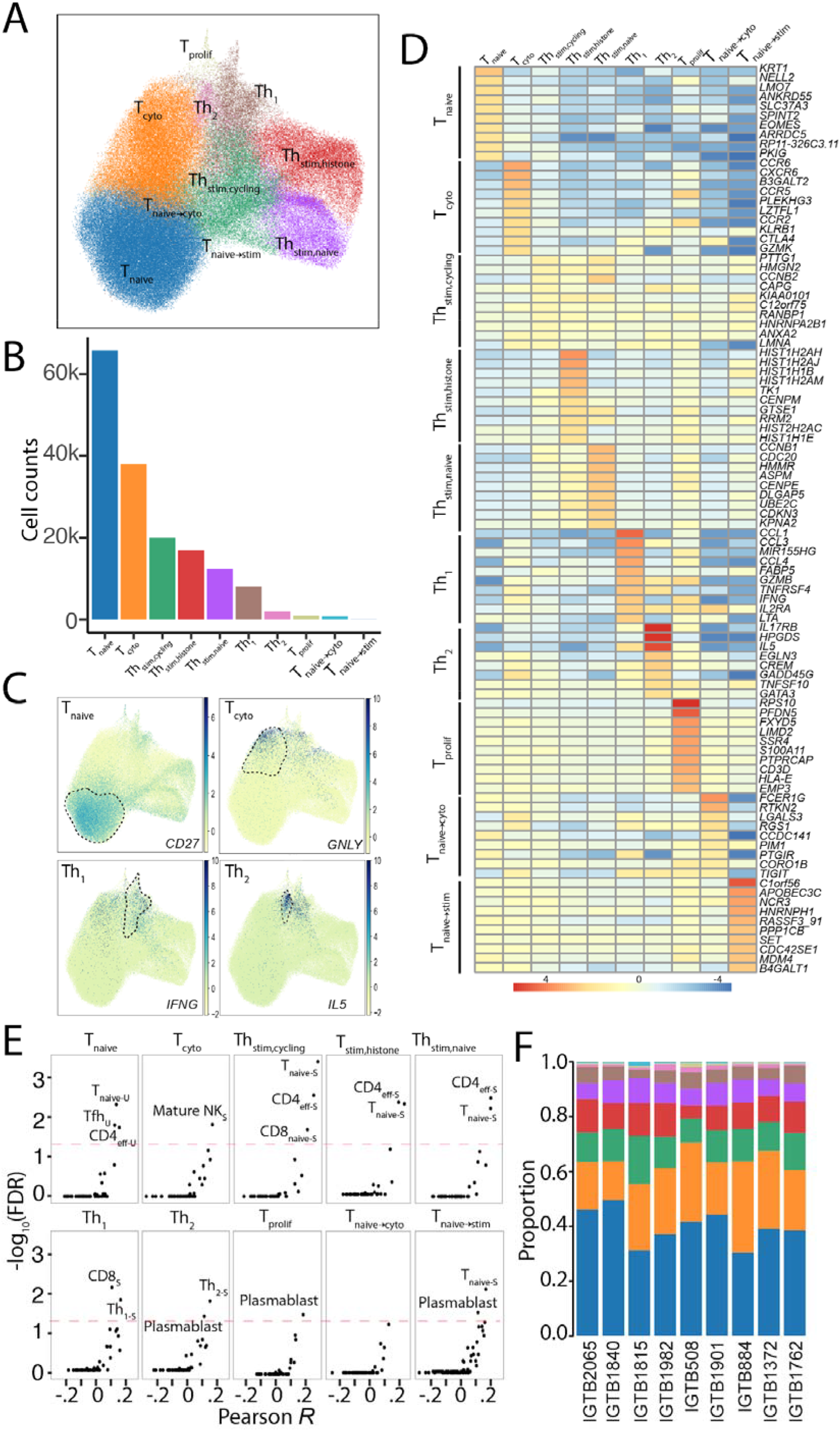
Heterogeneity of activated CD4^+^ T cells. **(A)** UMAP of activated CD4^+^ T cells. Each color represents an identified Leiden cluster. **(B)** Number of cells per cluster, colors correspond to the populations in (A). **(C)** Feature plots of normalized expression in UMAP coordinates of *CD27* (naive T; top left), *GNLY* (T_cyto_; top right), *IFNG* (Th_1_; bottom left), and *IL5* (Th_2_; bottom right). **(D)** Log fold-change (with respect to all other clusters) of top 10 positively differentially expressed (DE) genes (row) per cluster (column). **(E)** For each cluster, correlation of average log fold-change of DE genes to sorted bulk RNA sequencing transcriptomes (*22*) (x-axis) versus −log_10_(FDR) (y-axis). (**F**) Cluster proportions (y-axis) across nine donors (x-axis). Each color corresponds to the population in (A).

### Regulator perturbations drive T cell polarization

Regulators that control the activation and polarization of specific CD4^+^ T cell subsets have been mapped in mice and humans using either pooled CRISPR/Cas9 screens sorting for specific cell surface markers or RNA-interference (RNAi) followed by bulk transcriptomic profiling (*24*, *63*). These assays trade off perturbation throughput (low in RNAi, high in CRISPR/Cas9) and phenotypic resolution (low by cell sorting, high in bulk transcriptomic profiling). Here, we leverage the ability to link CRISPR perturbations to the transcriptomes of single cells using SLICE followed by mux-seq to enable high throughput (hundreds of loci) and high phenotypic resolution (transcriptome wide) mapping of regulators during human CD4^+^ activation and polarization. We first demonstrate the robustness and performance of our strategy by the following two quality control assessments. One, comparing cells expressing each knockout sgRNA (KO cells) to cells expressing the wild type sgRNAs (WT cells), the expression fold change for the targeted regulator was lower than random genes (FC_targeted_ = 0.56 *vs.* FC_random_ = 0; KS test; *P* < 2.26×10^−16^, Fig. 3A). Second, the transcriptomes of cells expressing sgRNAs targeting the same regulator or WT sgRNAs are more correlated on average (*R*_*KO*_ = 0.44, *R*_*WT*_ = 0.50) than cells expressing two random sgRNAs (*R*_*random*_= 0; KS test P < 2.2×10^−16^) (Fig. 3B). These two results suggest that sgRNAs targeting the same gene have similar downstream transcriptomic effects despite a modest (albeit statistically significant) change in overall fold change of the targeted genes.

**Figure 3.**
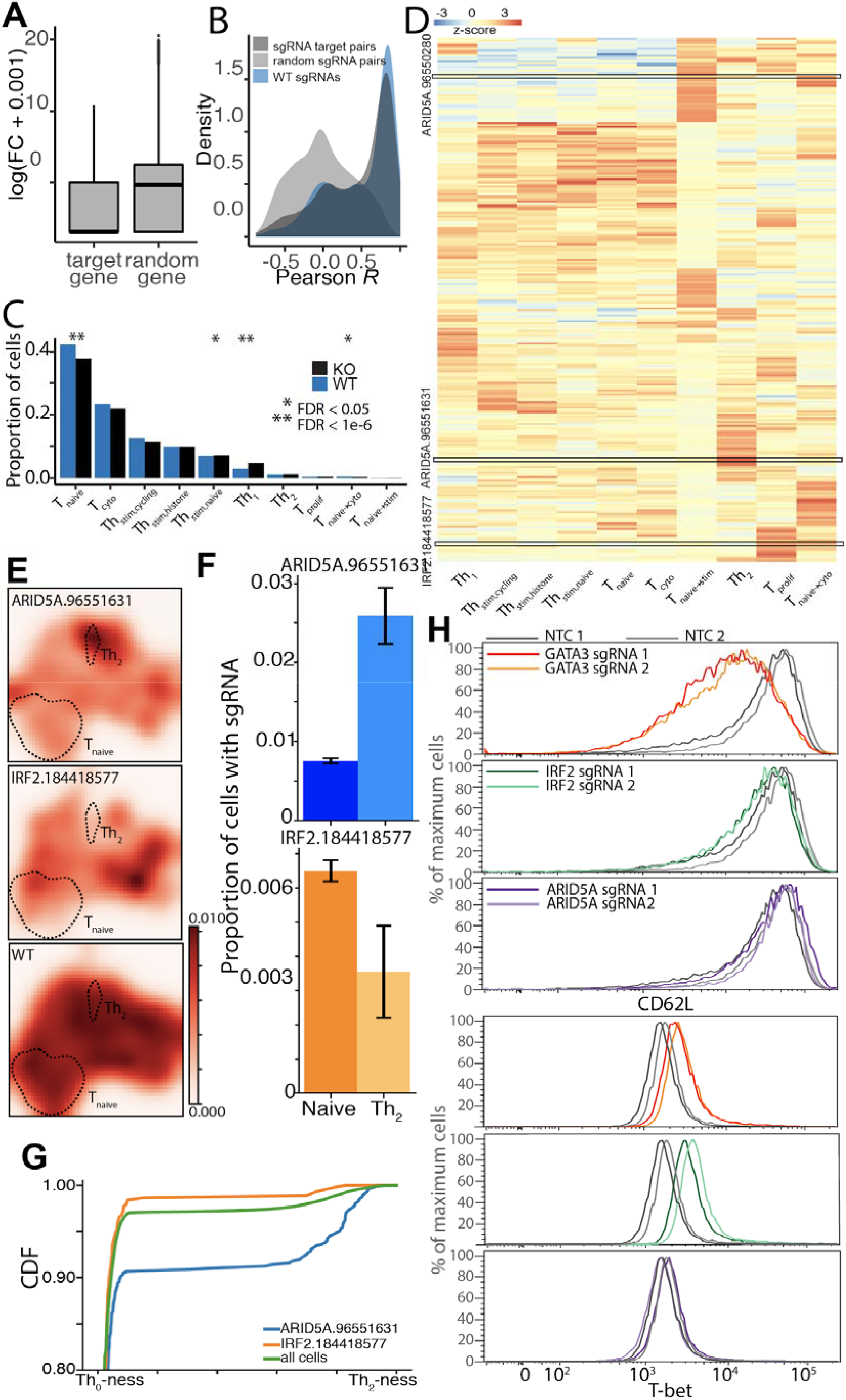
Regulator perturbations drive T cell polarization and maintenance. **(A)** Boxplot of fold change in expression of regulator targeted by sgRNA (left) and random gene (right) in cells expressing each sgRNA. **(B)** Distribution of transcriptomic correlations between cells expressing KO sgRNAs targeting the same gene (dark grey), WT sgRNAs (blue) and random sgRNA (light grey). **(C)** Proportion of KO (black) and WT (blue) cells per cluster. * indicates FDR < 0.05 and ** indicates FDR < 1e-6. **(D)** Clustered heatmap of sgRNA enrichment or depletion (z-score) across clusters. Red indicates a positive z-score and blue indicates a negative z-score. **(E)** Density of KO cells expressing sgRNA targeting *ARID5A ARID5A-*targeting (cutsite: chr2:96551631; top), *IRF2-*targeting (cutsite: chr4:184418577; middle), and WT (top) sgRNAs in UMAP space. **(F)** Proportion of cells expressing *ARID5A*-targeting (cutsite: chr2:96551631; top) or *IRF2-*targeting (cutsite: chr4:184418577; bottom) in T_naive_ and Th_2_ clusters. **(G)** Empirical cumulative distribution function (ECDF) of the estimated diffusion pseudotime of cells expressing sgRNA *IRF2-*targeting (cutsite: chr4:184418577; orange), cells expressing sgRNA *ARID5A*-targeting (cutsite: chr2:96551631; blue), and all cells (green). The shape of the ECDF reflects the enrichment of the guide along the pseudotime axis. **(H)** FACS validation. Distribution of cells expressing Th_2_ marker CD62L^+^ (top 3 panels) and Th_1_ marker T-bet^+^ (bottom 3 panels) electroporated with non-targeting control sgRNAs (grey), *GATA3-targeting* sgRNAs (red and orange), *IRF2*-targeting sgRNAs (light and dark green), or *ARID5A-*targeting sgRNAs (light and dark purple).

We next assessed the effects of KO sgRNAs on T cell states. Compared to WT cells, the proportion of KO cells is statistically enriched in activated or polarized subsets (e.g. Th_stim,naive_ and Th_1_) and depleted in the inactivated subsets (e.g. T_naive_ cells and transitional T_naive→cyto_; hypergeometric test, FDR < 0.05; Fig. 3C). In order to quantify the effect of each sgRNA on cell state, we computed the proportion of cells in each cluster that contained a particular sgRNA and identified those sgRNAs significantly enriched or depleted using a Z test (Fig. 3D; **Table 4**; **Methods**). Each cluster had on average 13 sgRNAs depleted (z-score < −1.5, P < 0.1) and 25 sgRNAs enriched (z-score > 1.5, P < 0.1; **Fig. S8**). For example, the sgRNA (cutsite: chr2:96551631, cutting efficiency: 0.47) targeting the RNA-binding protein AT-Rich Interactive Domain-Containing Protein 5A (*ARID5A*) was enriched in the Th_2_ cluster (z-score > 1.5, P = 9.0×10^−3^; Fig. 3E,**F**, top panels) and slightly depleted, although not statistically significantly, in the T_naive_ cluster (z-score <-0.4, P = 0.33). The second *ARID5A-*targeting sgRNA (cutsite: chr2:96550280, cutting efficiency: 0.097) showed consistent patterns of enrichment in T_naive_ and Th_2_ cells (T_naive_: z-score < −1.5, P < 0.1; Th_2_: z-score > 0.2, *P* = 0.35) (**Fig. S9**). In contrast, the sgRNA targeting interferon response factor 2 (*IRF2*; cutsite: chr4:184418577, cutting efficiency: 0.25) was depleted in Th_2_ cells (z-score < −1.5, P < 0.1; Fig. 3E, F, right panels). The second *IRF2-*targeting sgRNA (cutsite: chr4:184418667) had a cutting efficiency of 0.04 and was thus considered a WT sgRNA and is not enriched or depleted in any cluster (**Fig. S5**).

We next estimated the trajectory of polarization from T_naive_ to Th_2_ cells using diffusion pseudotime (DPT) and quantified the distribution of *ARID5A-*targeting (cutsite: chr2:96551631) and *IRF2-*targeting (cutsite: chr4:184418577) sgRNAs along the trajectory (**Methods**). The shape of the cumulative distribution function along the DPT is informative of enrichment or depletion of cells along the polarization trajectory. For example, if the cumulative percentage has an earlier increase, then the more likely a group of cells are to be naive and reside at an earlier “time-point”. Compared to all cells, cells containing the *IRF2-*targeting (cutsite: chr4:184418577) sgRNA are less likely to be Th_2_-like as shown by the 98.1% cumulative percentage at ~0.1 DPT (Fig. 3G), suggesting that *IRF2* could be important for the polarization of T_naive_ cells to Th_2_ cells or the maintenance of already polarized Th_2_ cells. In contrast, cells containing *ARID5A-*targeting (cutsite: chr2:96551631) sgRNA had a 90.6% cumulative percentage at ~0.1 DPT, exemplifying a greater enrichment at a later pseudotime (more Th_2_-like). This suggests that *ARID5A* may play a role in maintaining the T_naive_ cell phenotype, which is consistent with previous reports (*64*) (Fig. 3G).

To validate the *IRF2*- and *ARID5A-*targeting phenotypes, we used the Cas9 ribonucleoprotein (RNP) system to knockout *GATA3*, *ARID5A* and *IRF2* each with two sgRNAs and two non-targeting negative controls under general activation (anti-CD3/CD28) and Th_2_ (IL-4, anti-IFNG, anti-IL-12) polarization conditions (**Methods**). After two weeks of culturing, we used fluorescence-activated cell sorting (FACS) to sort for Th_2_ (CD62L^+^) and Th_1_ (T-bet^+^) cells and extracted DNA to assess cutting efficiency for each sample. In activated cells containing *IRF2*-targeting sgRNAs, the proportion of Th_2_s (CD62L^+^) was lower (3.81% and 2.02%) compared to non-targeting controls (3.46% and 7.71%) but higher compared to *GATA3-*targeting cells (1.26% and 0.892%; Fig. 3H, **S10**), consistent with the pooled screen results. Further, the proportion of Th_1_s (T-bet^+^) was higher in *IRF2-*targeting cells (10.5% and 8.02%) compared to non-targeting controls (0.87% and 1.93%) and *GATA3-*targeting cells (6.42% and 7.42%; Fig. 3H, **S10**). Interestingly, in Th_2_-polarized cells, there was not a change in the proportion of Th_2_ cells (**Fig. S11**). In contrast, in activated and Th_2_ polarized cells containing *ARID5A-*targeting sgRNAs, the proportion of Th_2_ cells are slightly higher (10.2% and 6.61%) and the proportion of Th_1_ cells remain unchanged (1.83% and 1.27%) (Fig. 3H, **S11**). These results recapitulate the pooled screen with *IRF2* acting as a positive regulator and *ARID5A* as a negative regulator of Th_s_s. Using a multiplexed pooled screening framework, we were able to harness the high resolution transcriptomic data to help elucidate and validate novel cell state regulators.

### Regulators interact to alter gene expression

Activation and polarization of T cells is known to involve the genetic interaction of regulators through direct physical cooperation, competition, and feedback and feedforward regulation of gene expression (*65*–*68*). For example, BATF and JUN physically interact to regulate transcription in dendritic cells (DCs), T cells, and B cells by jointly interacting with IRFs to bind compound-binding AP-1–IRF consensus elements (AICEs) (*69*). However, a map of genetic interactions between regulators that specify T cell function remain uncharted, primarily due to insufficient scalable methods to test for genetic interactions in primary T cells.

RNA interference or CRISPR perturbations followed by bulk RNA-seq allows us to study how genetic perturbations change gene expression on average across a population of cells. Detecting genetic interactions between regulators in this setting would require perturbing multiple regulators and observing non-additive changes in average expression, which is both experimentally and statistically intractable beyond a few regulators. By leveraging the ability to link genetic perturbations with their effects in many cells, we used SLICE followed by mux-seq to estimate the effects of CRISPR perturbations on the correlation between genes across cells to detect genetic interactions between regulators.

Specifically, we sought to map genetic interactions by identifying mutually mediating pairs of regulators, defined as one regulator modifying the correlation between another regulator with a downstream gene. As an example, if knocking out R_1_ modifies the correlation between R_2_ and G, then directionality is established as R_1_ *mediates* the effect of R_2_ on G (Fig. 4A) and vice versa. If R_1_ and R_2_ mutually mediate each other’s effects on G, we call R_1_ and R_2_ a genetic *interaction* and R_1_, R_2_, and G as a regulator (R) pair - gene triplet. To statistically detect mediation, we performed a likelihood ratio test between two linear mixed effect models, testing for the interaction term of R_1_ (presence of sgRNA targeting R_1_) and R_2,exp_ (R_2_ expression) (**Methods**). A significant change in correlation between expression of R_2,exp_ and a downstream gene suggests R_1_ mediates the effect of R_2_ on the downstream gene.

**Figure 4.**
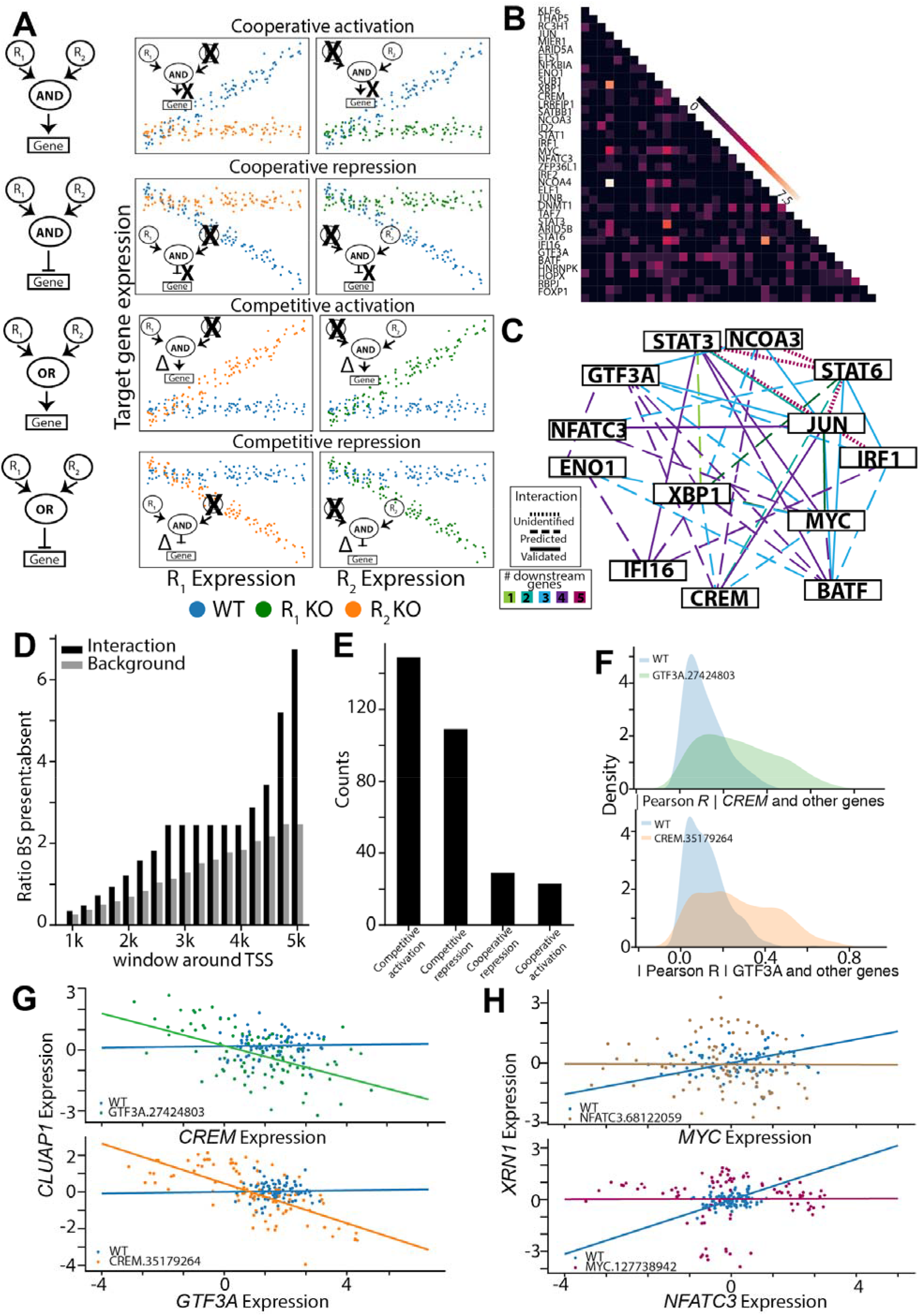
Perturbations and single cell analysis reveal transcription factor interactions. **(A)** Cartoon of detecting genetic interactions between regulators by comparing magnitude of correlation between KO and WT cells. In cooperative activation, magnitude of positive correlation decreases; in cooperative repression, magnitude of negative correlation decreases; in competitive activation, magnitude of positive correlation increases; in competitive repression, magnitude of negative correlation increases. **(B)** Number of genes downstream of each interacting regulator pair. **(C)** Network of interaction regulators known to affect T cell function. Solid line indicates known interactions; dashed indicates predicted interactions; dotted indicates known but undetected interactions. The colored edges indicate the number of downstream genes. **(D)** Ratio of identified target genes with both predicted binding sites to those without (y-axis) within a window size around the TSS (x-axis). **(E)** Distribution of regulator interactions found by subtype. **(F)** On top, distribution of magnitude of correlations between *CREM* and downstream genes for WT cells (blue) and *GTF3A* (cutsite: chr13:27424803) KO cells (green). On bottom, distribution of magnitude of correlations between *GTF3A* and downstream genes for WT cells (blue) and *CREM-*targeting (cutsite: chr10:35179264) KO cells (orange). **(G)** *CLUAP1* expression versus *CREM* expression in *GTF3A*-targeting (cutsite: chr13:27424803) KO cells in (top) and *GTF3A* expression in *CREM-*targeting (cutsite: chr10:35179264) KO cells (bottom). **(H)** *XRN1* expression versus *MYC* expression in *NFATC3*-targeting (cutsite: chr16:68122059) KO cells (top) and *NFATC3* expression in *MYC*-targeting (cutsite: chr8:127738942) KO cells (bottom), illustrating an example of cooperative activation. For **(G)** and **(H)**, trend lines reflects the coefficients fitted by the linear mixed effects model.

We identify four different types of regulator interactions: 1) cooperative activation, where regulators and the target gene are positively correlated in WT cells and uncorrelated in KO cells; 2) cooperative repression, where regulators and the target gene are negatively correlated in WT cells and uncorrelated in KO cells; 3) competitive activation, where regulators and the target gene are uncorrelated in WT cells and are positively correlated in KO cells; and 4) competitive repression, where regulators and target gene are uncorrelated in WT cells and are negatively correlated in KO cells (Fig. 4A).

We tested 37/140 of the most variably expressed regulators corresponding to total of 666 possible regulator pairs and 1,456,542 possible R pair - gene triplets (**Methods**). For each regulator, we first identified a set of downstream genes whose correlations with the regulator were affected when the regulator was perturbed using the same linear mixed effect model (Fig. 4A; **Methods**). For 33 of the 37 regulators (4 regulators each had a WT sgRNA), sgRNAs targeting the same regulator were more likely to identify the same downstream targets (*P* < 0.005, Mann-Whitney U test, **Fig. S12**) with similar changes in correlation (*R* = 0.31, *P* = 3.2×10^−78^; **Fig. S13**) compared to random (*R =* −0.004, *P* = 0.56).

To identify R pair - gene triplets, we intersected downstream genes for each regulator pair (FDR < 0.1) and tested for mutual mediation. We identified 310 R pair - gene triplets (FDR < 0.05) where the regulators mutually mediated each other’s effect on the downstream gene, comprised of 194 unique regulator pairs (Fig. 4B; **Table 5**). 24 of the regulator pairs identified were among the 48 regulator pairs previously known to interact (hypergeometric test, *P* < 0.005) (*70*). Combining all candidate genetic interactions reveals a core gene regulatory network in the functionalization of primary T cell (Fig. 4C) that suggests *JUN*, *MYC*, *XBP1*, and *STAT3/6* forming a central hub with many overlapping interacting partners. To validate the predicted regulator interactions and their downstream targets, we searched for transcription factor binding sites (TFBSs) of the 31 transcription factor (TF) pairs upstream and downstream of each predicted downstream gene’s transcription start site (TSS) that exist the Homer database (*71*) (**Methods**). We found a greater proportion of downstream gene TSSs containing TFBSs for both TFs compared to random sampling of TF pair - gene triplets (P < 0.05, Kolmogorov-Smirnov test across all windows, Fig. 4D).

We detected both known and novel interactions between key regulators for T cell functionalization. We identified two possible targets of the previously mentioned BATF-JUN interaction, *ATG14* and *TMEM204*. In addition, we identified an ETS1-STAT6 interaction, which is known to modulate cytokine responses in keratinocytes (*69*, *72*). While ETS1 has been shown to interact with numerous genes, in particular those in the STAT family, involved in the development and function of T cells (*73*), our result specifically suggests STAT6 as an interacting partner.

Amongst the candidate regulator interactions, we identified 23 pairs of cooperative activators, 29 pairs of competitive activators, 109 pairs of competitive repressors, and 149 pairs of competitive activators (Fig. 4E). In one example of competitive interaction, compared to WT cells, *GTF3A* is more correlated with genes in cells expressing the *CREM-*targeting (cutsite: chr10:35179264) sgRNA and *CREM* is more correlated with genes in cells expressing the *GTF3A-*targeting (cutsite: chr13:27424803) sgRNA (Fig. 4F). *CREM* and *GTF3A* is an example of a competitive repressor pair that regulates the expression of *CLUAP1*, where *CLUAP1* is negatively correlated with each regulator in KO cells but uncorrelated in WT cells (Fig. 4G). In another example, MYC and NFATC3 cooperatively interact to activate *XRN1* (Fig. 4H) where *XRN1* is positively correlated with each regulator in WT cells but not correlated in KO cells (FDR < 0.05). These results suggest that when a regulator is perturbed, downstream effects of other regulators become more prominent and this change can be harnessed to detect subtle interactions, often competitively activating interactions, between regulators.

### CRISPR perturbation modifies genetic effects on gene expression

While the contribution of interindividual variability to the composition, expression and activation of CD4^+^ T cells *ex vivo* has been described by us (*18*, *19*) and others (*20*, *74*–*79*), little is known about the interindividual variability in CRISPR perturbed cells. Using a linear mixed model, we analyzed cells expressing each sgRNA to identify 125 genes whose expression were variable between individuals (interindividual genes, FDR < 0.2) across 79 sgRNAs (Fig. 5A, B; **Table 7**; **Methods**).

**Figure 5.**
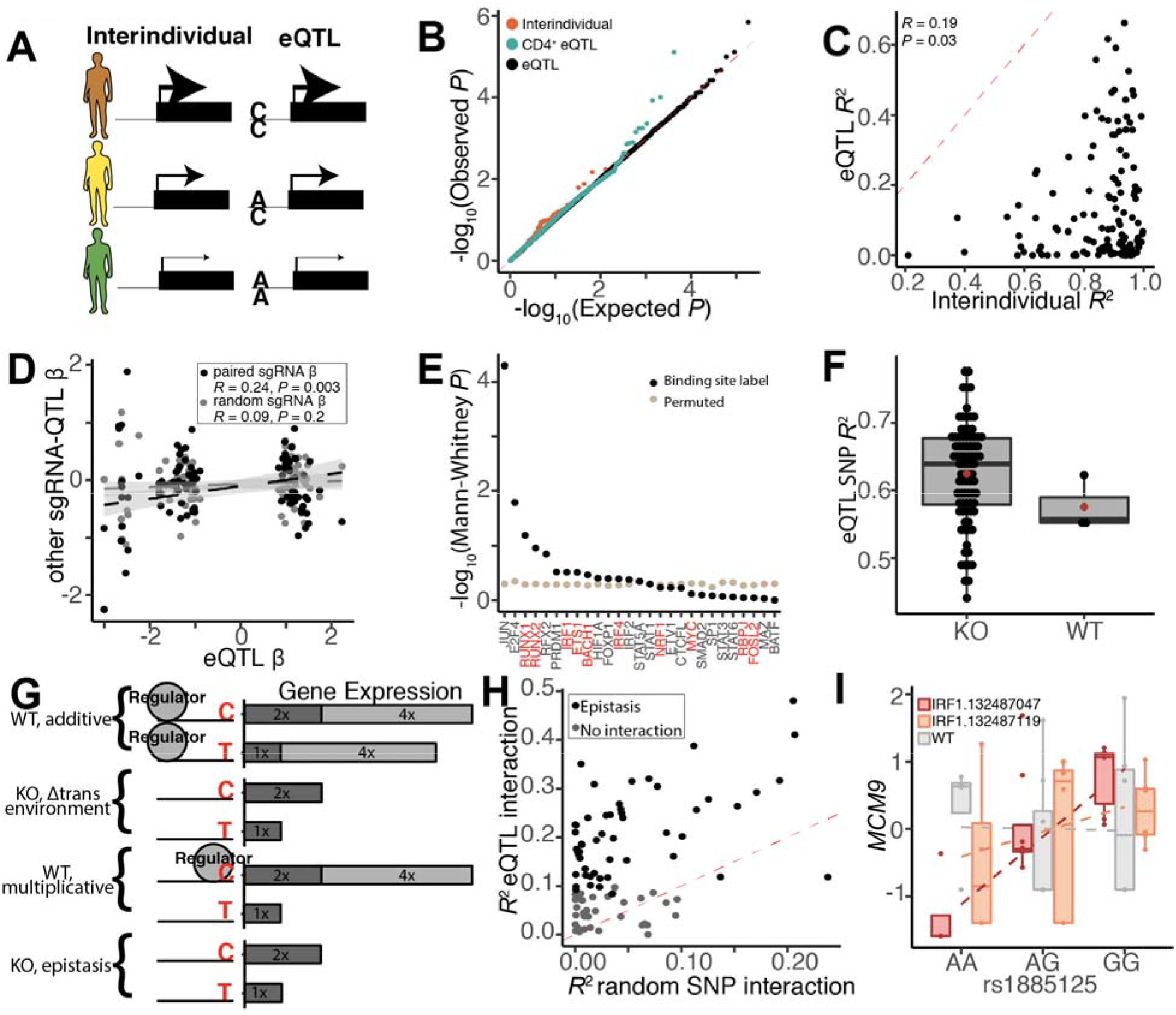
CRISPR perturbation modifies genetic effects on gene expression. **(A)** Cartoon of interindividual variation of gene expression (left) and possible genetic causes due to a SNP located in the cis regulatory region (right). Size of the arrow corresponding to the amount of gene expression for each donor. **(B)** eQTL QQ-plot. Each point represents an eGene empirical *P* value across sgRNAs (black), and those that are also interindividual genes (orange) and previously identified CD4^+^ eGenes (teal). X-axis: expected *P* values. Y-axis: observed *P* values. Red dashed line is null. **(C)** Scatter plot of variance explained by interindividual (x-axis) and genetic (y-axis) variation, per eGene. **(D)** Scatter plot of eQTL effect sizes between pairs of sgRNAs targeting the same gene (black) or not (gray). **(E)** −log10 Mann-Whitney *P* (y-axis) of observing the ranked order of genes harboring binding sites for each regulator. Black is observed and tan is for permuted binding sites, and regulators in red have an eQTLs. **(F)** Genetic variance explained of significant eQTLs (y-axis) in KO and WT cells (x-axis). **(G)** Cartoon depicting genetic ablation impact on the effects of a donor with a C and another with a T allele on gene expression. If the regulator has an additive effect on gene expression in WT (first cartoon), then regulator absence changes the *trans* environment (second cartoon). If regulator interacts with a *cis* regulatory element to have a multiplicative effect (third cartoon), then regulator absence changes the effect of the SNP (fourth cartoon). **(H)** Variance explained by *cis* x *trans* interaction for our eQTLs (y-axis) and a random SNP interaction (x-axis). In black are significant interactions. **(I)** Normalized *MCM9* expression (y-axis) is subsetted by donor genotype (x-axis) at rs1885125 in *IRF1-*targeting (chr5:132487047, red), *IRF1*-targeting (chr5:132487119, orange), and WT (grey) cells.

However, interindividual variability can be attributed to genetic, environmental and technical confounding effects. To identify the genetic contribution, we performed an expression quantitative trait loci (eQTL) analysis using a linear mixed model that includes a genetic covariate term (Fig. 5A; **Methods**). Because of the limited number of samples, we significantly reduced the multiple testing burden by only testing for SNPs +/− 100 kb around a TSS with a minor allele frequency > 0.4 and only highly expressed genes per sgRNA (on average 1,891 genes). We found a total of 88 eQTLs (in *cis*) across cells expressing all sgRNAs (permutation FDR < 0.2) corresponding to 84 genes (eGene) (Fig. 5B). To assess the robustness of these results, we performed two quality check analyses. First, genes previously reported to have an eQTL in activated CD4^+^ T cells (*18*) and with significant interindividual variability were more statistically significant than those that were not (Fig. 5B). Second, the variance explained by genetics was correlated with and expectedly less than the variance due to interindividual variation amongst eGenes (*R* = 0.19, *P* = 0.03, Fig. 5C).

We next assessed eQTLs detected in KO versus WT cells. Overall, eQTL effect sizes were more correlated between cells expressing sgRNAs targeting the same regulator (*R* = 0.24, *P* = 0.003) than between cells expressing a random pair of sgRNAs (*R* = 0.09, *P* = 0.2), suggesting genetic effects specific to each knockout (Fig 5D). For the regulators that exist in the Homer database (*71*), we found that genes harboring binding sites for 10 regulators are more likely to be eQTLs (as indicated by more significant *P*) (Fig. 5E). On average, genetics explained 64% and 56% of the variability in KO and WT cells respectively (Fig. 5F). Finally, we found eQTLs were more likely to be detected in KO cells (22% of KO *vs.* 14% of WT sgRNAs; hypergeometric *P* = 0) (**Fig. S14**). These results support that *in vitro* genetic perturbations by CRISPR/Cas9 can uncover effects of natural genetic variation undetectable in unperturbed cells.

The increased power to detect genetic effects on gene expression only in KO cells could be due to a change in the *trans* environment or regulator-genetic interactions (*cis* x *trans* epistatic interaction) (Fig. 5G). If the activity of a regulator has an additive effect on gene expression, then ablating the regulator will decrease the overall variance of gene expression thereby increasing the genetic contribution to gene expression (**Fig. 5F**, **S15-17**). Supporting this model, 80 out of the 88 eQTLs had lower standard errors in KO cells compared to WT cells (**Fig. S16**). If the activity of a regulator multiplicatively interacts with genetic variants (epistasis), then ablating the regulator should change the genetic effect on gene expression. To identify instances of epistasis, we fit a linear mixed model testing for *cis* (SNP) x *trans* (sgRNA presence or absence) interactions (Fig. 5G). We found statistical evidence for epistasis for 48 out of 88 eQTLs (FDR < 0.05), where the sgRNA is more likely to interact with the eQTL than a random SNP (Fig. 5H). These results suggest that CRISPR/Cas9 ablation of a regulator can uncover both additive and epistatic effects from standing genetic variation on gene expression.

As an illustrative example, we found evidence for an epistatic interaction between *IRF1*-targeting (cutsite: chr5:132487047) sgRNA and genetic variant (rs1885125) to regulate *MCM9* expression (Fig. 5H). *IRF1* (also known as interferon regulatory factor 1) has previously been shown to regulate T cell activation (*80*–*82*), particularly driving Th_1_ polarization (*83*). Two independent epigenetic analyses suggest IRF1 binding at the *MCM9* promoter (**Fig. S18**). First, in K562 cells, there is an IRF1 ChIP-seq peak 142 bp upstream from rs1885125, containing four SNPs in LD (D’ > 0.97). Second, the Homer database (*71*) predicted an IRF1 binding site 595 bp upstream from rs1885125 using an IRF1 ChIP-seq in peripheral blood mononuclear cells, which is flanked by rs4946371 (D’ > 0.98). These results support the interaction between a genetic variant in a *cis* regulatory element of *MCM9* and the *trans* factor, IRF1, to account for 47% of *MCM9* expression variability. All together, these results suggest that CRISPR perturbations can uncover interindividual variation in gene expression and in some cases, epistatically interact with natural standing variation to modulate the variability of gene expression.

## Discussions

CD4^+^ T cells serve diverse roles in the adaptive immune system by dynamically responding to extracellular signals in their microenvironment. While the gene regulatory networks governing these responses have been extensively studied in mice, the topology and parameters of these networks, including how they vary across individuals, have not been mapped in humans. To address these gaps, we present the first large scale, multiplexed single cell RNA-seq study of activated primary CD4^+^ T cells isolated from 9 donors across CRISPR perturbations targeting hundreds of candidate regulators.

Activated CD4^+^ T cells are heterogeneous, capturing cell states reminiscent of canonical helper subtypes (e.g. Th_1_, Th_2_, etc), cytotoxic phenotypes, and broad activation or cell cycle. We find that cells expressing WT sgRNAs are more likely to be T_naive_ cells while those expressing KO sgRNAs are more likely to promote polarization into a differentiated state (e.g. Th_1/2_ cells), exemplified by the identification *ARID5A* as a negative regulator of Th_2_ polarization. We expect that the application of our approach to cells differentiated or polarized under specific conditions (e.g. toward a Th_2_ phenotype) rather than broad activation (e.g. anti-CD3/CD28 activation) can further refine the mapping of gene regulatory programs that control T cell differentiation, polarization and maintenance.

The identification of gene gene interactions is experimentally and combinatorially challenging in primary cells. By exploiting coexpression patterns between single cells, we devised a new approach based on differential correlation analysis to detect interactions between pairs of regulator during T cell activation and polarization. Using this approach, we reconstructed a gene regulatory network for T cell activation that includes known interactions (e.g. *JUN, MYC, XBP1, STAT*) and previously unreported interactions such as between *ETS1* and *STAT6*, which may be involved in the propagation of T cell cytokine signaling. By increasing the number of cells profiled using multiplexed workflows (*30*, *84*) and the number of genetic perturbations through higher multiplicity of infection (*85*), future integration of genome engineering and single cell transcriptomics would allow for refined mapping and causal reconstruction (*86*) of gene regulatory networks in specific low frequency T cell subsets (e.g., Th_1/2_, T_naive_).(*86*)

Although epistatic interactions involving naturally segregating variants have been identified in model organisms, there has been been limited success in identifying these interactions using observational studies in humans due to limited power (*87*–*89*). Our genetic-multiplexed approach allowed us to identify genes that are interindividual variable in CRISPR perturbed primary human cells and in some cases, pinpoint the genetic variants that likely mediate the variability. Akin to reducing the *trans* contribution of gene expression through *in vitro* perturbations(*18*–*20*, *90*) or computational adjustments (*91*), we provide evidence of decreased gene expression variance in CRISPR perturbed cells thus increasing the ability to detect *cis* genetic effects. Surprisingly, we also found that some CRISPR perturbations can modify the effects of genetic variants on gene expression epistatically reminiscent of gene by environment effects detected by *in vitro* perturbation of cells (*18*–*20*, *90*). Thus, a comprehensive perturbative-QTL analysis using CRISPR/Cas9 is a compelling alternative strategy to large observational studies for mapping genetic interactions that involve standing genetic variants in primary human cells.

Our work provides the first view into the heterogeneity of activated CD4^+^ T cells at the single cell resolution across pooled CRISPR perturbations and individuals. We identify candidate regulators of T cell polarization and two classes of genetic interactions. By harnessing natural and CRISPR genome engineering, we can begin to efficiently dissect gene regulatory networks and identify genetic interactions in primary human cells.

## Acknowledgments

We would like to thank all Immvar participants. We also thank all members of the Ye lab for laboratory discussions.

## Author contributions

A.L. performed the SLICE experiment, sgRNA amplicon preparation, and paragon sequencing preparation. D.L. performed the Th2 validation experiment. R.E.G. performed all alignments, demuxlet, cell identification, cell state identification, sgRNA enrichment and depletion, interindividual, and genetic variation analyses. M.C.K. identified the sgRNAs in each cell and performed the DPT and regulator interaction analyses. M.S. performed eQTL simulations. R.E.G., M.C.K., and C.J.Y. wrote the manuscript.

## References

1. L. Guglani, S. A. Khader, Th17 cytokines in mucosal immunity and inflammation. Curr. Opin. HIV AIDS. 5, 120–127 (2010).

2. A. Kimura, T. Kishimoto, Th17 cells in inflammation. Int. Immunopharmacol. 11, 319–322 (2011).

3. S. Leung et al., The cytokine milieu in the interplay of pathogenic Th1/Th17 cells and regulatory T cells in autoimmune disease. Cell. Mol. Immunol. 7, 182–189 (2010).

4. D. Liu et al., IL-25 attenuates rheumatoid arthritis through suppression of Th17 immune responses in an IL-13-dependent manner. Sci. Rep. 6, 36002 (2016).

5. D. A. Rao, T Cells That Help B Cells in Chronically Inflamed Tissues. Front. Immunol. 9, 1924 (2018).

6. L. Petersone et al., T Cell/B Cell Collaboration and Autoimmunity: An Intimate Relationship. Front. Immunol. 9, 1941 (2018).

7. S. Kuchen et al., Essential role of IL-21 in B cell activation, expansion, and plasma cell generation during CD4+ T cell-B cell collaboration. J. Immunol. 179, 5886–5896 (2007).

8. S. M. Kerfoot et al., Germinal center B cell and T follicular helper cell development initiates in the interfollicular zone. Immunity. 34, 947–960 (2011).

9. M. Goodman, T cell regulation of polyclonal B cell responsiveness. III. Overt T helper and latent T suppressor activities from distinct subpopulations of unstimulated splenic T cells. Journal of Experimental Medicine. 153 (1981), pp. 844–856.

10. R. J. Hodes, T helper cell-b cell interaction: the roles of direct Th-B cell contact and cell-free mediators. Semin. Immunol. 1, 33–42 (1989).

11. D. C. Parker, T cell-dependent B cell activation. Annu. Rev. Immunol. 11, 331–360 (1993).

12. F. P. Legoux et al., CD4+ T Cell Tolerance to Tissue-Restricted Self Antigens Is Mediated by Antigen-Specific Regulatory T Cells Rather Than Deletion. Immunity. 43, 896–908 (2015).

13. A. K. Abbas, J. Lohr, B. Knoechel, V. Nagabhushanam, T cell tolerance and autoimmunity. Autoimmunity Reviews. 3 (2004), pp. 471–475.

14. A. Takeuchi, T. Saito, CD4 CTL, a Cytotoxic Subset of CD4+ T Cells, Their Differentiation and Function. Front. Immunol. 8, 194 (2017).

15. J. van Bergen et al., Phenotypic and functional characterization of CD4 T cells expressing killer Ig-like receptors. J. Immunol. 173, 6719–6726 (2004).

16. J. J. Zaunders et al., Identification of circulating antigen-specific CD4+ T lymphocytes with a CCR5+, cytotoxic phenotype in an HIV-1 long-term nonprogressor and in CMV infection. Blood. 103, 2238–2247 (2004).

17. X. Zhou et al., Instability of the transcription factor Foxp3 leads to the generation of pathogenic memory T cells in vivo. Nat. Immunol. 10, 1000–1007 (2009).

18. R. E. Gate et al., Genetic determinants of co-accessible chromatin regions in activated T cells across humans. Nat. Genet. 50, 1140–1150 (2018).

19. C. J. Ye et al., Intersection of population variation and autoimmunity genetics in human T cell activation. Science. 345, 1254665 (2014).

20. T. Raj et al., Polarization of the effects of autoimmune and neurodegenerative risk alleles in leukocytes. Science. 344, 519–523 (2014).

21. S. Kasela et al., Pathogenic implications for autoimmune mechanisms derived by comparative eQTL analysis of CD4+ versus CD8+ T cells. PLoS Genet. 13, e1006643 (2017).

22. D. Calderon et al., Landscape of stimulation-responsive chromatin across diverse human immune cells (2018),, doi:10.1101/409722.

23. P. Zhou et al., In vivo discovery of immunotherapy targets in the tumour microenvironment. Nature. 506, 52–57 (2014).

24. N. Yosef et al., Dynamic regulatory network controlling TH17 cell differentiation. Nature. 496, 461–468 (2013).

25. J. Henriksson et al., Genome-wide CRISPR Screens in T Helper Cells Reveal Pervasive Crosstalk between Activation and Differentiation. Cell. 176 (2019), pp. 882–896.e18.

26. B. Adamson et al., A Multiplexed Single-Cell CRISPR Screening Platform Enables Systematic Dissection of the Unfolded Protein Response. Cell. 167, 1867–1882.e21 (2016).

27. A. Dixit et al., Perturb-Seq: Dissecting Molecular Circuits with Scalable Single-Cell RNA Profiling of Pooled Genetic Screens. Cell. 167, 1853–1866.e17 (2016).

28. P. Datlinger et al., Pooled CRISPR screening with single-cell transcriptome readout. Nat. Methods. 14, 297–301 (2017).

29. E. Shifrut et al., Genome-wide CRISPR Screens in Primary Human T Cells Reveal Key Regulators of Immune Function. Cell. 175, 1958–1971.e15 (2018).

30. H. M. Kang et al., Multiplexed droplet single-cell RNA-sequencing using natural genetic variation. Nat. Biotechnol. 36, 89–94 (2018).

31. K. Schumann et al., Generation of knock-in primary human T cells using Cas9 ribonucleoproteins. Proceedings of the National Academy of Sciences. 112, 10437–10442 (2015).

32. A. J. Hill et al., On the design of CRISPR-based single-cell molecular screens. Nat. Methods. 15, 271–274 (2018).

33. K. K.-H. Farh et al., Genetic and epigenetic fine mapping of causal autoimmune disease variants. Nature. 518, 337–343 (2015).

34. V. Traag, L. Waltman, N. J. van Eck, From Louvain to Leiden: guaranteeing well-connected communities (2018), (available at http://arxiv.org/abs/1810.08473).

35. J. J. Campbell et al., CCR7 expression and memory T cell diversity in humans. J. Immunol. 166, 877–884 (2001).

36. T. Worbs, R. Förster, A key role for CCR7 in establishing central and peripheral tolerance. Trends Immunol. 28, 274–280 (2007).

37. A. Kanhere et al., T-bet and GATA3 orchestrate Th1 and Th2 differentiation through lineage-specific targeting of distal regulatory elements. Nature Communications. 3 (2012),, doi:10.1038/ncomms2260.

38. J. Zhu, H. Yamane, J. Cote-Sierra, L. Guo, W. E. Paul, GATA-3 promotes Th2 responses through three different mechanisms: induction of Th2 cytokine production, selective growth of Th2 cells and inhibition of Th1 cell-specific factors. Cell Research. 16 (2006), pp. 3–10.

39. A. O’Garra, L. Gabryšová, Transcription Factors Directing Th2 Differentiation: Gata-3 Plays a Dominant Role. The Journal of Immunology. 196 (2016), pp. 4423–4425.

40. B. Jakiela, W. Szczeklik, H. Plutecka, Increased production of IL-5 and dominant Th2-type response in airways of Churg–Strauss syndrome patients (2012) (available at https://academic.oup.com/rheumatology/article-abstract/51/10/1887/1821359).

41. S. Greenfeder, S. P. Umland, F. M. Cuss, R. W. Chapman, R. W. Egan, Th2 cytokines and asthma. The role of interleukin-5 in allergic eosinophilic disease. Respir. Res. 2, 71–79 (2001).

42. T. R. Mosmann, R. L. Coffman, TH1 and TH2 cells: different patterns of lymphokine secretion lead to different functional properties. Annu. Rev. Immunol. 7, 145–173 (1989).

43. G. Seumois et al., Transcriptional Profiling of Th2 Cells Identifies Pathogenic Features Associated with Asthma. J. Immunol. 197, 655–664 (2016).

44. P. Angkasekwinai et al., Interleukin 25 promotes the initiation of proallergic type 2 responses. J. Exp. Med. 204, 1509–1517 (2007).

45. N. M. Weathington et al., IL-4 Induces IL17Rb Gene Transcription in Monocytic Cells with Coordinate Autocrine IL-25 Signaling. Am. J. Respir. Cell Mol. Biol. 57, 346–354 (2017).

46. L. M. Bradley, D. K. Dalton, M. Croft, A direct role for IFN-gamma in regulation of Th1 cell development. J. Immunol. 157, 1350–1358 (1996).

47. R. B. Smeltz, J. Chen, R. Ehrhardt, E. M. Shevach, Role of IFN-in Th1 Differentiation: IFN-Regulates IL-18R Expression by Preventing the Negative Effects of IL-4 and by Inducing/Maintaining IL-12 Receptor 2 Expression. The Journal of Immunology. 168 (2002), pp. 6165–6172.

48. E. L. Filbert, M. Le Borgne, J. Lin, J. E. Heuser, A. S. Shaw, Stathmin regulates microtubule dynamics and microtubule organizing center polarization in activated T cells. J. Immunol. 188, 5421–5427 (2012).

49. J. A. Best et al., Transcriptional insights into the CD8(+) T cell response to infection and memory T cell formation. Nat. Immunol. 14, 404–412 (2013).

50. J. E. Noll et al., PTTG1 expression is associated with hyperproliferative disease and poor prognosis in multiple myeloma. J. Hematol. Oncol. 8, 106 (2015).

51. D. Wu et al., Impact of PTTG1 downregulation on cell proliferation, cell cycle and cell invasion of osteosarcoma and related molecular mechanisms. Zhonghua bing li xue za zhi= Chinese journal of pathology. 43, 695–698 (2014).

52. S. Fan, X. Li, L. Tie, Y. Pan, X. Li, KIAA0101 is associated with human renal cell carcinoma proliferation and migration induced by erythropoietin. Oncotarget. 7, 13520–13537 (2016).

53. R.-H. Yuan et al., Overexpression of KIAA0101 predicts high stage, early tumor recurrence, and poor prognosis of hepatocellular carcinoma. Clin. Cancer Res. 13, 5368–5376 (2007).

54. M. Jain, L. Zhang, E. E. Patterson, E. Kebebew, KIAA0101 is overexpressed, and promotes growth and invasion in adrenal cancer. PLoS One. 6, e26866 (2011).

55. C. Chevaleyre et al., The Tumor Antigen Cyclin B1 Hosts Multiple CD4 T Cell Epitopes Differently Recognized by Pre-Existing Naive and Memory Cells in Both Healthy and Cancer Donors. J. Immunol. 195, 1891–1901 (2015).

56. P. L. Brazee et al., FXYD5 Is an Essential Mediator of the Inflammatory Response during Lung Injury. Front. Immunol. 8, 623 (2017).

57. I. L. Gotliv, FXYD5: Na/K-ATPase Regulator in Health and Disease. Frontiers in Cell and Developmental Biology. 4 (2016),, doi:10.3389/fcell.2016.00026.

58. I. Lubarski-Gotliv, C. Asher, L. A. Dada, H. Garty, FXYD5 Protein Has a Pro-inflammatory Role in Epithelial Cells. J. Biol. Chem. 291, 11072–11082 (2016).

59. H. Peng et al., LIMD2 Is a Small LIM-Only Protein Overexpressed in Metastatic Lesions That Regulates Cell Motility and Tumor Progression by Directly Binding to and Activating the Integrin-Linked Kinase. Cancer Research. 74 (2014), pp. 1390–1403.

60. F. Wang et al., LIMD2 targeted by miR⍰34a promotes the proliferation and invasion of non⍰small cell lung cancer cells. Mol. Med. Rep. 18, 4760–4766 (2018).

61. D. Wang et al., Prefoldin 1 promotes EMT and lung cancer progression by suppressing cyclin A expression. Oncogene. 36, 885–898 (2017).

62. F. A. Wolf, F. Hamey, M. Plass, J. Solana, J. S. Dahlin, Graph abstraction reconciles clustering with trajectory inference through a topology preserving map of single cells. bioRxiv (2018) (available at https://www.biorxiv.org/content/10.1101/208819v2.abstract).

63. M. Ciofani et al., A validated regulatory network for Th17 cell specification. Cell. 151, 289–303 (2012).

64. K. Masuda et al., Arid5a regulates naive CD4+ T cell fate through selective stabilization of Stat3 mRNA. J. Exp. Med. 213, 605–619 (2016).

65. T. Ravasi et al., An atlas of combinatorial transcriptional regulation in mouse and man. Cell. 140, 744–752 (2010).

66. E. V. R. Hao Yuan Kueh, Regulatory gene network circuits underlying T-cell development from multipotent progenitors. Wiley Interdiscip. Rev. Syst. Biol. Med. 4, 79 (2012).

67. C. Hernandez-Munain, J. L. Roberts, M. S. Krangel, Cooperation among Multiple Transcription Factors Is Required for Access to Minimal T-Cell Receptor α-Enhancer Chromatin In Vivo. Mol. Cell. Biol. 18, 3223 (1998).

68. A. Kröger, IRFs as competing pioneers in T-cell differentiation. Cellular and Molecular Immunology. 14, 649 (2017).

69. T. L. Murphy, R. Tussiwand, K. M. Murphy, Specificity through cooperation: BATF–IRF interactions control immune-regulatory networks. Nat. Rev. Immunol. 13, 499 (2013).

70. C. von Mering et al., STRING: known and predicted protein-protein associations, integrated and transferred across organisms. Nucleic Acids Res. 33, D433–7 (2005).

71. S. Heinz et al., Simple Combinations of Lineage-Determining Transcription Factors Prime cis-Regulatory Elements Required for Macrophage and B Cell Identities. Mol. Cell. 38, 576–589 (2010).

72. J. Travagli, M. Letourneur, J. Bertoglio, J. Pierre, STAT6 and Ets-1 Form a Stable Complex That Modulates Socs-1 Expression by Interleukin-4 in Keratinocytes. J. Biol. Chem. 279, 35183–35192 (2004).

73. H. V. Nguyen et al., The Ets-1 transcription factor is required for Stat1-mediated T-bet expression and IgG2a class switching in mouse B cells. Blood. 119, 4174–4181 (2012).

74. J. D. Storey et al., Gene-Expression Variation Within and Among Human Populations. Am. J. Hum. Genet. 80, 502–509 (2007).

75. T. Lappalainen et al., Transcriptome and genome sequencing uncovers functional variation in humans. Nature. 501 (2013), pp. 506–511.

76. L. Chen et al., Genetic Drivers of Epigenetic and Transcriptional Variation in Human Immune Cells. Cell. 167, 1398–1414.e24 (2016).

77. J. F. Degner et al., DNase I sensitivity QTLs are a major determinant of human expression variation. Nature. 482, 390–394 (2012).

78. W. Zhang et al., A global transcriptional network connecting noncoding mutations to changes in tumor gene expression. Nat. Genet. 50, 613–620 (2018).

79. F. Grubert et al., Genetic Control of Chromatin States in Humans Involves Local and Distal Chromosomal Interactions. Cell. 162, 1051–1065 (2015).

80. S. Giang, A. La Cava, IRF1 and BATF: key drivers of type 1 regulatory T-cell differentiation. Cell. Mol. Immunol. 14, 652–654 (2017).

81. T. Ohteki et al., The Transcription Factor Interferon Regulatory Factor 1 (IRF-1) Is Important during the Maturation of Natural Killer 1.1 T Cell Receptor–α/β (NK1 T) Cells, Natural Killer Cells, and Intestinal Intraepithelial T Cells. The Journal of Experimental Medicine. 187 (1998), pp. 967–972.

82. J. D. Brien et al., Interferon Regulatory Factor-1 (IRF-1) Shapes Both Innate and CD8+ T Cell Immune Responses against West Nile Virus Infection. PLoS Pathog. 7, e1002230 (2011).

83. S.-I. Kano et al., The contribution of transcription factor IRF1 to the interferon-gamma-interleukin 12 signaling axis and TH1 versus TH-17 differentiation of CD4+ T cells. Nat. Immunol. 9, 34–41 (2008).

84. C. S. McGinnis et al., MULTI-seq: Scalable sample multiplexing for single-cell RNA sequencing using lipid-tagged indices,, doi:10.1101/387241.

85. M. Gasperini et al., A Genome-wide Framework for Mapping Gene Regulation via Cellular Genetic Screens. Cell. 176, 1516 (2019).

86. K. Yang, A. Katcoff, C. Uhler, in International Conference on Machine Learning (2018), pp. 5541–5550.

87. A. H. Y. Tong, Global Mapping of the Yeast Genetic Interaction Network. Science. 303, 808–813 (2004).

88. A. P. Davierwala et al., The synthetic genetic interaction spectrum of essential genes. Nat. Genet. 37, 1147–1152 (2005).

89. B. Lehner, C. Crombie, J. Tischler, A. Fortunato, A. G. Fraser, Systematic mapping of genetic interactions in Caenorhabditis elegans identifies common modifiers of diverse signaling pathways. Nat. Genet. 38, 896–903 (2006).

90. M. N. Lee et al., Common Genetic Variants Modulate Pathogen-Sensing Responses in Human Dendritic Cells. Science. 343 (2014), pp. 1246980–1246980.

91. D. Lee, A. Cheng, N. Lawlor, M. Bolisetty, D. Ucar, Detection of correlated hidden factors from single cell transcriptomes using Iteratively Adjusted-SVA (IA-SVA). Sci. Rep. 8, 17040 (2018).

