## Supplementary material for "Mapping gene regulatory networks of primary CD4^+^ T cells using single-cell genomics and genome engineering": Methods

**Study subjects and genotyping**

Our samples were enrolled in PhenoGenetic study (age 18 to 56, average 29.9), as part of the Immvar cohort(*20*), which were recruited in the Greater Boston Area. Each donor gave written consent to participate and were healthy, without any history of inflammatory disease, autoimmune disease, chronic metabolic disorders or chronic infectious disorders. We genotyped 56 caucasian samples on the OmniExpressExome54 chip, and excluded 2080 SNPs with a call rate <90% (0.22% of total), 1521 SNPs with Hardy Weinberg *P* < 0.0001 (0.16%) and 259,860 SNPs with MAF < 0.01 (27.04%) out of the total 960,919 SNPs profiled. The Michigan Imputation Server was used to impute these genotypes with the Haplotype Reference Consortium Panel Version r1.1. After genotype imputation had 5,324,560 SNPs, which were then subsetted for our nine donors.

**Regulator target identification**

Our library contained targeted 140 regulators (transcription factors and RNA-binding proteins) with 2 sgRNAs each. Each regulator was unbiasedly chosen using gene expression and accessibility data from activated CD4^+^ T cells in 95 and 105 healthy donors(*18*). To get the highly expressed regulators using RNA-seq data, we performed a TMM normalization and took the upper quartile of highly expressed genes and subsetted those that were regulators. To get the regulators with highly accessible binding sites using ATAC-seq data, we enriched for all binding sites on the HOMER database(*71*) in activated accessible chromatin regions. We took the union of the highly expressed regulators and accessible binding sites, for a total of 140 regulators (**Fig. 1B**).

**CROP-seq library generation**

The backbone plasmid used to clone the CROP-Seq library was CROPseq-Guide-Puro(*28*), purchased from Addgene (Addgene. Plasmid #86708). We used two sgRNAs oligo sequences from the Brunello library(*88*) for each of our chosen 140 regulators. Oligos for the sgRNA library were purchased from Integrated DNA Technologies (IDT) and cloned into the CROPseq plasmid backbone using the methods described by Datlinger et al. (2017)(*28*). Lentivirus was produced using the UCSF ViraCore.

**SLICE experiment and sequencing**

Primary human CD4^+^ T cells were isolated from peripheral blood mononuclear cells (PBMCs) by magnetic negative selection using the EasySep Human CD4^+^ T Cell Isolation Kit (STEMCELL, Cat #17952). Cells were cultured in X-Vivo media, consisting of X-Vivo15 medium (Lonza, Cat #04- 418Q) with 5% Fetal Calf Serum, 50mM 2-mercaptoethanol, and 10mM N-Acetyl L-Cysteine. On the day of isolation (Day 1), cells were rested in media without stimulation for 24 hours. The day after isolation (Day 2), cells were stimulated with ImmunoCult Human CD3/CD28 T Cell Activator (STEMCELL, Cat #10971) and IL-2 at 50U/mL. 24 hours post stimulation (Day 3), 1 uL of lentivirus was added directly to cultured T cells and gently mixed. Following 24 hours (Day 4), cells were collected, pelleted, and washed in PBS twice. Then, cells were resuspended in Lonza electroporation buffer P3 (Lonza, Cat #V4XP-3032). Cas9 protein (MacroLab, Berkeley, 40mM stock) was added to the cell suspension at a 1:10 v/v ratio. Cells were transferred to a 96 well electroporation cuvette plate (Lonza, cat #VVPA-1002) for nucleofection using the Lonza Nucleofector 96-well Shuttle System and pulse code EH115 (Lonza, cat #VVPA-1002). Immediately after electroporation, pre-warmed media was added to each electroporation well, and 96-well plate was placed at 37 degrees for 20 minutes. Cells were then transferred to culture vessels in X-Vivo media containing 50U/mL IL-2 at 1e6 cells /mL in appropriate tissue culture vessels. Two days later, 1.5ug/mL Puromycin was added in culture media for selection. Cells were expanded every two days, adding fresh media with IL-2 at 50U/mL. Cells were maintained at a cell density of 1e6 cells /mL. On the final day (Day 13) of the experiment, cells from each of the nine donors were counted using Vi-CELL XR and pooled at equal numbers to obtain a final 180,000 cells in 60 uL of PBS (**Fig. 1A**). The pooled cells were then processed by UCSF Institute for Human Genetics (IHG) Genomics Core using 16 wells of 10X Chromium Single Cell v2 (PN-120237), as per manufacturer’s protocol, with each well being separately index. The final library was sequenced on two lanes on the Nova-seq for a total of 6.7B reads.

**10x transcriptome alignment**

Each 10x well was separately aligned to our personalized reference, which contained the hg19 transcriptome and our 280 sgRNA sequences using cellranger “count” function. Our reference sgRNA sequences contained the U6 plasmid promoter on the 5’ and sgRNA scaffold on the 3’, such that our sgRNA reference sequences were as follows:

TATGCTTACCGTAACTTGAAAGTATTTCGATTTCTTGGCTTTATATATCTTGTGGAAAGGACGAAACACCG - 20 bp gRNA - GTTTTAGAGCTAGAAATAGCAAGTTAAAATAAGGCTAGTCCGTTATCAACTTGAAAAAGTGGCACCGAGT. Using the cellranger “aggr” function, we aggregated all 16 wells into one combined dataset.

**Demuxlet**

Per well, demuxlet(*30*) was run on all 737,280 raw error corrected barcodes with 10x cellranger bam file, using a 1% genotype error rate, 0.5 alpha, 255 minimum mapping quality, 0 minimum distance to tail. We combined all 16 demuxlet runs, then we used the “BEST” column to identify droplet multiplet and the “SNG.1ST” column to identify the donor of origin.

**sgRNA amplicon sequencing and analysis**

For the sgRNA amplicon sequencing, donors were separately amplified and barcoded using a two-step PCR protocol. First, each donor was divided into 8 PCR reactions with 0.1ng template of cDNA. Each 25mL reaction consisted of 1.25mL P5 forward primer, 1.25mL Nextera Read 2 reverse primer, priming to the U6 promoter to enrich for guides, 12.5mL NEBNext Ultra II Q5 Master Mix (NEB, cat #M0544L), 0.1ng template, and water to 25mL. The PCR cycling conditions were: 3 minutes at 98C, followed by 10 s at 98C, 10 s at 62C, 25 s at 72C, for 10 cycles, and a final 2 minute extension at 72C. After the PCR, all reactions were pooled for each donor and purified using Agencourt AMPure XP SPRI beads (cat #A63881) per the manufacturer’s protocol. Next, 1mL was taken from each purified PCR product to go into a second PCR for sequencing index. Each reaction included 1mL of PCR product, 12.5 mL NEBNext Ultra II Q5 Master Mix (NEB, cat #M0544L), 1.25mL P5 forward primer, 1.25mL Illumina i7 primer, and water to 25mL. The PCR cycling conditions were: 3 minutes at 98C, followed by 10 s at 98C, 10 s at 62C, 25 s at 72C, for 10 cycles, and a final 2 minute extension at 72C. After the PCR, all reactions were SPRI purified and quantified using the Qubit dsDNA high sensitivity assay kit (Thermo Fisher Scientific, cat# Q32854) and run on a gel to confirm 500bp size.

The sgRNA amplicon sequencing library was sequenced paired-end on one lane HiSeq 4000, resulting in 171 million mapped paired-end reads. This library was similarly aligned to our personalized reference containing our 280 sgRNA sequences using 10x “count” function. We generated a read count matrix of 320,708 barcodes x 280 sgRNA matrix.

**sgRNA identification**

In order to identify the sgRNAs present in each cell, we performed a series of binomial test of enrichment for the top 4 sgRNAs present each of our 320,708 cells. Let n be the total number of reads mapping to any sgRNA in a given cell. Let c1, c2, c3, c4 be the number of reads of the top four sgRNA in a given cell, with c1 representing the sgRNA with the most reads. We performed a binomial test with c1 successes from trails = n - c2 - c3 - c4, with 1/280 probability of success. This test for the enrichment of the sgRNA with c1 counts, disregarding the other three of the top four sgRNAs. We performed a similar test with the guide with c2 read count, performing a binomial test of c2 successes from trails = n - c1 - c3 - c4 trials, with 1/280 probability of success. The binomial tests statistics were bonferroni corrected *P* < 0.05(**Table S2**).

**Cell identification**

To identify our cell barcodes, we used our combined transcriptomic dataset and our combined demuxlet output, as previously mentioned. We filtered out cell-containing droplets that contained less than 10 SNPs from our demuxlet results, totaling 320,708 cells. 10 SNPs was the threshold at which the number of cell-containing barcodes and then number of cell-containing barcodes with at least 1000 UMIs no longer increased (**Fig. S1**, **S2**, **S3**). Cells were filtered for demuxlet identified singlets, which resulted in a 24% doublet rate, and cells where we could not identify the sgRNA (25%), resulting in 164,624 cell (**Fig. 1C**, **Table S2**).

**sgRNA cutting efficiencies**

Genomic DNA was isolated from cell pellets using the Promega Wizard Genomic DNA Purification Kit (cat #A1120). Amplification was performed as described by Paragon User Guide (CleanPlex Custom NGS Panel). Briefly, gDNA template was added to a multiplex PCR reaction with 2 uL of 5X Paragon PCR Mix, 2 uL of 5X Primer Pool, gDNA and Nuclease-Free Water. Each sgRNA amplicon was designed to be ~200bp, centering around the cutsite(**Table S1**). The PCR cycling conditions were: 10 minutes at 95C, followed by 15 cycles of 15 seconds at 98C and 5 minutes at 60C. Following the PCR, amplified DNA was purified using Magnetic Beads from Paragon and subjected to a digestion reaction (CP Reagent Buffer, CP Digestion Reagent) to remove nonspecific PCR products. A post-digestion purification was performed, followed by a second PCR reaction to amplify and index libraries. Second PCR reaction contained 5X Second PCR Reaction Master Mix, purified DNA from previous step, and i5/i7 Indexed PCR Primers for Illumina. The PCR cycling conditions were: 10 minutes at 95C, followed by 10 cycles of 15 seconds at 98C and 75 seconds at 60C. Finally, DNA was purified using magnetic beads from Paragon and subjected to next-generation sequencing.

**Paragon sgRNA amplicon DNA sequencing analysis**

Per donor, our sgRNA DNA sequencing was aligned to hg19 using bwa -mem(*89*). Using the “mpileup” function from the samtools suite of tools(*90*, *91*) we estimated the number of indels and reads per basepair +/-200bp around each sgRNA cutsite(**Table S3**). Per sgRNA and per donor, we estimated indel frequency as the maximal number reads with indels +/- 5 target cutsite divided by total reads covering a cutsite. Cutting efficiencs for each sgRNA were then estimated as the indel frequency x the proportion in the original sgRNA library.

**sgRNA WTs**

Using our sgRNA amplicon DNA sequencing we identified WT sgRNAs by calculating a z-score for each sgRNA. Let i be a sgRNA, and c is the cutting efficiency of sgRNA_i_, and p is the proportion of cells with sgRNA_i_, then WT sgRNAs were identified as z-scores of p / c with *P* < 0.05. In total we identified 14 WT sgRNAs, which have the maximum cutting efficiencies was < 5% with at least 484 cells. To estimate WT proportions (**Fig. 2E**), we calculated a hypergeometric *P* comparing the number of WT cells and KO cells per cluster as a function of all total WT and KO cells.

**Single cell normalization**

We normalized our 164,623 cells using the scanpy(*92*) suite of tools. Working with our 32,739 genes x 164,1623 cell matrix, we calculated the percentage of mito contamination, filtered out genes using “filter_genes” with options “min_counts=1” and then normalized using “normalize_per_cell”, which normalizes each gene cell by the total counts for that cell. We further filtered our gene list for variable genes, which we identified by subsetting the cells from one well, then calling “filter_genes_dispersion” with options “min_mean=0.0125, max_mean=3, min_disp=0.5”. Using only one well safe guarded us from variable genes due to well to well batch effects. Subsetting our gene x cell matrix for our 2,189 variables genes, we re-normalized our cells for total sequencing, log transformed, and then regressed out mitochondrial contamination (as previously calculated), the 10x well, and total UMIs. Finally, we scaled the data to have mean = 0 and variance = 1. To reduce the data down two dimensions for visualization, we ran UMAP(*93*) through scanpy(*92*) using our 2,189 variable genes across our 164,623 cells(**Fig. 3A**).

**Leiden clustering**

We performed unbiased cluster detection on our 164,623 cells using our 2,189 variable genes, using the scanpy(*92*) suite of tools. First, we calculated a neighborhood graph(*93*), second, performed leiden clustering(*34*) at 0.68 resolution. We compared our leiden clusters at 0.5, 0.6, 0.68, 0.75, and 1 resolutions and found that our clusters called at a 0.68 resolution were qualitatively the most similar to the gene expression patterns of T cell subtype markers (Th_2_: IL5, Th_1_:IFNG, naive T: CD27).

**Differential analysis**

For cluster differential expression, we used our 2,189 variable gene x 164,1623 cell normalized matrix (as previously described), which was subsetted by donor. Using the “FindAllMarkers” function from the Seurat package(*94*) using options “min.pct=0, logfc.threshold=0, min.cells.gene=0, min.diff.pct=0, return.thresh=1” we calculated the log fold-change for all 2,189 genes per donor. We then performed a meta analysis to estimate a meta *P* per gene across all nine donors using the metap package in (*95*) and we averaged the log fold-changes across all nine donors to get an average log fold-change.

For sgRNA differential expression, we again started with our 2,189 variable gene x 164,1623 cell normalized matrix, which was subsetted by donor and by sgRNA. Using the “FindMarkers” function from the Seurat package(*94*) using options “min.pct=0, logfc.threshold=0, min.cells.gene=0, min.diff.pct=0, return.thresh=1” we compared cells containing each KO sgRNA to our WT cells, per donor. Again, we calculated a meta *P* per gene and log fold-changes were averaged across the nine donors per gene per sgRNA.

**Comparison to sorted, bulk T cell subtypes**

Using the RNA-seq dataset from Calderon et al. 2018(*96*), we averaged the gene expressions across all donors for each cell type and to normalize it, we log transformed the expression counts, subtracted out the median count per sample, and then standardized by the gene (mean = 0, variance = 1). For each cell type we took the top 300 most highly expressed genes, and then correlated those 300 genes to their respective log average fold-changes from the cluster differential expression analysis. We calculated a *P*, which was then FDR adjusted.

**Th_2_ validation experiment**

PBMCs were sourced from anonymized female caucasian donors and were purified from whole blood by Ficoll gradient. Cells were frozen in 10% DMSO in FBS in a cryostorage vessel for one day at -80°C before being moved into a liquid nitrogen tank. Frozen PBMCs were quickly thawed in a 37°C water bath and slowly diluted with RPMI1640(Sigma, R0883) supplemented with 10% FBS(HI-FBS; Invitrogen, catalog 10438026), 1 mM GlutaMAX(Invitrogen; catalog 35050061), 100 U/ml penicillin and 100 mg/ml streptomycin(Invitrogen; catalog 15140122). Cells were pelleted at 300 xg for 5 minutes before being washed with SepMate Buffer(Stem Cell; catalog 20144) for naïve CD4^+^ isolation.

Naïve CD4^+^ T cells were isolated using a EasySep™ Human Naïve CD4^+^ T Cell Isolation Kit II(catalog 17555) according to the manufacturer’s protocol. Harvested naïve CD4 cells(1-3x10^6^) were plated on a 24 well plate with 1 ml of supplemented RPMI1640 with 50 ng/ml IL-2(R&D; catalog 202-IL-010) and 25 ul/ml of ImmunoCult™ Human CD3/CD28 T Cell Activator(StemCell Technologies, catalog 10971) for 24 hours. For differentiation modulation, activated T cells were split into 16 groups(5x10^4^-2x10^5^) per donor (n=7) for 8 guides and two polarizing conditions(Th_2_, activated CD4^+^). For RNP electroporation, 4 ul of 160 uM of tracr RNA (Dharmacon, catalog U-002005-50) was incubated with 4 ul of 160uM sgRNA (Dharmacon) for 30 minutes at 37°C. Following the annealing of sgRNAs, 8 ul of 40 uM Cas9-NLS(MacroLab, Berkeley, 40 uM stock) was added to each sgRNA mixture for 15 minutes at 37°C. 3 ul of each complete RNP was added to a 96 U-bottom plate(Genesee Scientific; catalog 25-221) alongside 1 ul of ssODN Alt-R Cas9 Electroporation Enhancer(IDT: catalog 1075916; 100 uM). Cells were then pelleted at 90 x g for 5 minutes at room temperature. Cells were resuspended in 20 ul of P3 buffer(Lonza; catalog V4SP-3096) and transferred to the aliquoted RNP mixtures. 20 ul of the cell-RNP mixture was added to a 96-well electroporation plate(Lonza, V4SP-3096) and electroporated on a 4D nucleofector system(Lonza) with program EH-115. Cells were quickly rescued by adding 100 ul of supplemented RPMI with 50 ng/ml IL-2 dropwise to electroporated cells. Cells were incubated for 10 minutes in a 37°C incubator with 5% CO_2_ before being transferred into a 96 well U-bottom plate and brought up to a total volume of 200 ul with 50 ng/ml IL-2 for activated CD4^+^ groups and 10 ng/ml IL-4(R&D, catalog 204-IL-010), 2 ug/ml anti-IL-12 antibody(R&D, catalog MAB219-500), and 2 ug/ml anti-IFN-G antibody(R&D, catalog MAB285-500) for Th_2_ and incubated for 24 hours. Media with appropriate cytokines were refreshed every 2-3 days and cell density was adjusted to 1x10^6^ cells/well with every media change. After a total of 14 days, cells were harvested for flow cytometry.

For maintenance modulation, naïve T cells were isolated as described. Cells were incubated with supplemented RPMI1640 with 50 ng/ml IL-2 and 25 ul/ml Immunocult for 72 hours in a 96 well U-bottom plate in 200 ul. Cells were then split into a Th_2_ and activated CD4^+^ conditions as described and cultured for 1 week, with cytokine supplemented media every 2-3 days. Cells were then split into 16 groups and electroporated as previously described for each guide. Cells were harvested 7 days after electroporation for flow cytometry.

For flow cytometry, cells were stimulated with PMA and ionomycin with Brefeldin A (Leukocyte activation cocktail with BD Golgiplug; BD Bioscience; catalog 550583) for five hours before staining. Cells were then washed twice with 1% BSA in PBS by pelleting cells at 300 x g for 5 minutes at 4°C and resuspended in staining buffer (Biolegend cell staining buffer; Biolegend; catalog 420201) with 5 ul of Trustain FCX(Biolegend; 422302) and incubated for 5 minutes on ice. Cells were then stained with 5 ul of each extracellular antibody(CD4-FITC, BV785-CD62L, AF700-CD127, BV711-CRTH2, BV510-CCR5; Biolegend) for 30 minutes on ice in the dark in a total reaction volume of 100 ul. After staining, cells were washed twice with 1% BSA in PBS. Cells were fixed and permeabilized using a eBioscience FOXP3/Transcriptional factor staining kit(eBioscience, 00-5523-00) per manufacturer’s protocol. Permeabilized cells were stained with 5 ul of each intracellular marker(BV421-GATA-3, APC-T-bet, BV605-IL-4, PE-IFN-G; Biolegend) and incubated for 30 minutes on ice in the dark for a total reaction volume of 100 ul. Cells were washed twice with 1% BSA in PBS and resuspended in 200 ul of PBS before flow analysis. BD LSRII(Parnassus Flow Core, Grover) was used for flow acquisition and FlowJo 9 was used for analysis.

For TIDE validation, 10^4^ cells were placed into 50 ul of Quickextract(Lucigen, catalog QE09050) and vortexed for 15 seconds. The cell solution was then incubated at 65C for 6 minutes and vortexed for another 15 seconds. The cell solution was then placed into a heat block at 98C for 2 minutes. The extracted gDNA was then stored at -20°C until amplification. Primers for targeted genes were designed to create a 700 bp amplicon, starting from 350 bp upstream of the cut site for the guide. Primers were designed using Primer-Blast(NCBI). Sequencing primers designed to be 200 bp upstream of the cut site were designed using the same tool. For amplification, 1 ul of gDNA solution was amplified using KAPA Hotstart HIFI Readymix (Kapa Biosystems, catalog KK2602). Generated amplicons were sent for sequencing with designed sequencing primers. Analysis of provided chromatograms were done using a TIDE web tool(<https://tide.deskgen.com>) and cutting efficiency was determined.

**Target gene expression**

We created a pseudobulked matrix, per sgRNA, per donor, and per cluster for a 32,739 genes x 17,845 samples, which was then filtered for our targeted regulators genes. Per sample, let i be each of our 140 targetted regulators, n is the total counts per sample, and r is counts for regulator_i_. Now, let w counts for regulator_i_ WT sample and m is total counts for the WT sample, then the fold-change_i_ = (r_i_ / c ) / (w_i_ / m ). As background, we randomly sampled a regulator for each sample and performed the same calculations (**Fig. 3A**).

**Correlation of sgRNAs**

We pseudobulked by sgRNA, by donor, and by cluster, for a total of 2,189 variables gene 17,845 samples x matrix. We normalized our matrix using using a log2 transformed median normalization and then standardized across a gene(mean=0, variance=1). For every sample, we averaged the gene expression across our 9 donors and clusters, and then correlated the normalized, averaged transcriptome for every pair of sgRNAs targeting the same regulator. As background, we correlatsed 280 unpaired sgRNA (**Fig. 3B**).

**sgRNA cluster enrichment/depletion**

We subsetted our cells by those that contained KO sgRNAs and calculated the sgRNA proportion per cluster. Then, we calculated a z-score per cluster across all KO sgRNAs. KO sgRNAs that had a z-score > 1.5 were considered enriched in that cluster and those that had z-scores < -1.5 were considered depleted in that cluster (**Fig. 3C**). To visualize sgRNA enrichment and depletion, the UMAP space (as previously described) was partitioned into a grid of 50x50 rectangular pixels, and the density of cells with a specific guide was computed in each rectangle. Gaussian blurring with sigma 3 was applied to the UMAP density image (**Fig. 3D**).

**Lineage trajectory**

First, we estimated diffusion pseudotime from the naive T cells to the Th_2_ cluster using the scanpy(*92*) implementation of Haghverdi et al. 2016(*97*). Using our normalized 2,189 gene x 164,623 cell matrix (as previously described), we filtered out cells that were not in either the naive T cell or the Th_2_ cluster. PCA, neighborhood construction (500 neighbors, 40 PCs), and UMAP were re-run prior to applying the diffusion pseudotime (DPT) algorithm, all with default parameters. DPT is a random-walk-based distance that is computed based on simple Euclidian distances in the 'diffusion map space'. The diffusion map is a nonlinear method for recovering the low-dimensional structure underlying high-dimensional observations(*97*). This algorithm assigns a single number to each cell, corresponding to the “time” that each cell has passed from a root cell. An empirical cumulative density plot was created using these estimated pseudotimes to detect and visualize distinct DPT profiles of cells containing different sgRNA.

Second, we used the scanpy implementation of the partition-based graph abstraction (PAGA) algorithm to quantify the connectivity of all of our cell clusters, approximating the overall cellular trajectory manifold. The default parameters for the PAGA algorithm was used, and connectivity > 0.3 was used for visualization.

**Regulator - Regulator interaction model**

We created ten technical replicates of pseudobulks by sgRNA and by donors, for a 2,189 variable gene x 5,040 sample matrix, which was normalized by estimating the proportion of total reads per gene for each sample, which was then multiplied by the median total reads across samples. Then, the data was log transformed and standardized(mean=0, variance=1). Out of the 140 regulators considered in our study, 37 were included when we selected for genes with highly variable expression using scanpy(*92*). To identify potential downstream genes for each regulator, we used a linear mixed model exp(G) ~ exp(regulator) + regulator_KO + exp(regulator):regulator_KO + intercept and donor as a random effect, testing for the addition of the interaction term exp(regulator):regulator_KO via a likelihood ratio test. Once potential downstream genes were identified for each regulator, pairs of regulators were formed and candidate regulator-pair, gene triplets were formed based on the intersection of potential downstream genes of those two regulators (regulator1 and regulator2). For every candidate regulator1, regulator2, G triplet, we test the interaction term in the following linear mixed models: exp(G) ~ exp(regulator1) + regulator2_KO + exp(regulator1):regulator2_KO + intercept, and exp(G) ~ exp(regulator2) + regulator1_KO + exp(regulator2):regulator1_KO + intercept, both using donor as a random effect. If the interaction terms are significant in both of the linear mixed models via likelihood ratio test, we call this regulator1, regulator2, G triplet an *interaction*.

**Regulator - Regulator interaction binding site validation**

Of the 37 regulators considered for the regulator interaction analysis, 18 had bindings sites in the HOMER database. 31 interactions of regulator-pair gene triplets were identified for this validation, where both regulators are in the HOMER database. For each regulator-pair gene triplet in this set, we searched for binding sites of regulator1 and regulator2 upstream and downstream of gene G’s TSS at various window length by using HOMER’s annotatePeaks.pl program(*71*). Window lengths were ranged from +/- 1 to 5 kbps in intervals of 250 bps. For each window, the ratio of number of interactions with both bindings sites in the window to the number of interactions without both binding sites in the window was calculated. The background was generated by taking all variable genes and randomly assigning pairs of those 18 regulators and applying the same procedure of looking for both binding sites in within the window around the TSS of each gene.

**Interindividual variation analysis**

We created two technical replicates of pseudobulks by sgRNA and by donors, for a 32,739 gene x 5,040 sample matrix. We filtered for genes with at least 10 counts and then further filtered for genes with a SNP +/- 100kb from the TSS with a minor allele frequency > 0.4, for a final 2,095 tested genes. We normalized each sgRNA separately, such that we subsetted our matrix 2,095 genes x 18 samples, where we calculated the percentage of total reads per gene and multiplied by the median total counts for all samples. Then, we log transformed the data and standardized it (mean=0, variance=1). For each sgRNA we also created a covariate file, containing donor and cutting efficiency, which was also standardized (mean=0, variance=1).

To test for interindividual variation, we used a linear mixed model, using the “Lme4” package(*98*). Per sgRNA we tested two models, our alternative model: expr ~ cutting efficiency + (1|donor), and our null model: expr ~ cutting efficiency. As Storey, et al. 2007(*74*) noted, including donor as a random effect properly accounts for donor variation, rather than a fixed effect. We calculated a likelihood ratio *P* to determine if donor was significant. Using the package “r.squaredGLMM” function from the MuMIn R package we calculated the R^2^ for each model, where the variance explained due to interindividual variation was calculated as the difference between the alternative and the null (R^2^ interindividual variation = R^2^ alternative - R^2^ null).

We calculated empirical *P*-values per gene per sgRNA. We permuted the donors, while maintaining donor pairs, 1000 times per gene, for a total of 527,282,000 permutations. To calculate empirical *P*-values first, we filtered interindividual variation associations by those that converged, and then filtered our permuted *P*-values for duplicate *P*-values. The former filtering step was performed because we wanted to reduce our multiple testing burden and therefore did not want to include tests that did not converge. The latter filtering step was performed because if permuting the donors caused that specific model to not converge, and if that occured multiple times, then that could inflate out statistics. Using the remaining, unique list of *P*-values we calculated empirical *P-*values using “empPvals” function from the q value package(*99*) with the option “pool=T”. Finally, we FDR adjusted our empirical *P-*values to determine significant interindividual associations.

**eQTL analysis**

Using the normalized expression matrices and covariate files for each sgRNA from our interindividual variation analysis we associated each gene to a genetic variation. As previously mentioned, we only tested variants that had a minor allele frequency > 0.4 and were +/-100kb around a TSS of a tested gene.

To detect eQTLs, we fit a linear mixed model, per sgRNA, where we fit two models, alternative: expr ~ SNP + cutting efficiency + (1|donor), and our null: expr ~ cutting efficiency + (1|donor). We performed a likelihood ratio and determine if the genetic statistical significance. Similarly to our interindividual association analysis, we used the function “r.squaredGLMM” function from the MuMIn R package to calculate R^2^ for each model. The variance explained due to genetics SNP R^2^ = R^2^ alternative - R^2^ null.

To calculate empirical *P*-values, per sgRNA, we permuted our genotypes 1000 times per gene - SNP test. We performed 1000 permutations tests per sgRNA per gene, in total we performed 301,020,000 permutations. To calculate well calibrated empirical *P*-values, per sgRNA we pooled all *P*-values and calculated empirical *P*-values using the “empPvals” function from the q value package(*99*) with the “pool=T” option, and then FDR adjusted. To test for sgRNA specific eQTLs, we recalculated FDRs per gene across all 268 sgRNAs, filtering for sgRNAs that did not test that gene.

**Binding site enrichment in eGenes**

27 out of our 140 regulators are in the the Homer database(*71*), therefore we parsed each gene for our 27 regulator binding site +/-100 kb around it’s TSS. Per tested regulator, we ranked our eQTL associations by *R*^2^ and compared our ranked list of genes that did and did not contain a binding site using a Mann-Whitney test. For each gene - regulator pair, we permuted the labels of genes that did and did and not have the binding site 100 times, calculating a Mann-Whitney *P* per test, taking the average of the permuted *P* (**Fig. 5I**).

**Bootstrapping variance explained**

To overcome our unbalanced sample sizes between KO and WT sgRNAs, we performed sampled each KO eQTL (with replacement) to the depth of our WT eQTLs (three eQTLs), 100 times. Per bootstrap, we estamed the mean and standard deviation of the variance explained across the 3 sampled KO eQTLs (**Fig. 5K**).

**Epistasis analysis**

Using the normalized expression matrices and covariate files for each sgRNA from our interindividual variation and eQTL analysis we performed an interaction test for the 88 eQTLs. For each eQTL we compared the KO sgRNA to a randomly sampled WT sgRNA condition. We fit two models, alternative: expr ~ SNP*sgRNA + cutting efficiency + (1|donor), and our null: expr ~ SNP + sgRNA + cutting efficiency + (1|donor). We performed a likelihood ratio and determine interaction statistical significance. We used the function “r.squaredGLMM” function from the MuMIn R package to calculate R^2^ for each model. The variance explained due to the interaction is R^2^ = R^2^ alternative - R^2^ null.

**eQTL standard error simulations**

Using an effect size of 0.5 and minor allele frequency of 0.5, we first simulated WT expression as the sum of the genetic effect, regulator effect, and an independent noise term (WT=g*beta + tf_expr + noise) and KO expression as the genetic effect and an independent noise term (KO=g*beta + noise). Next, we performed a linear regression on the WT and KO conditions and calculated the effect sizes, standard errors, and p-values for 1000 iterations.
