## Supplemental Figures for "Mapping gene regulatory networks of primary CD4^+^ T cells using single-cell genomics and genome engineering"

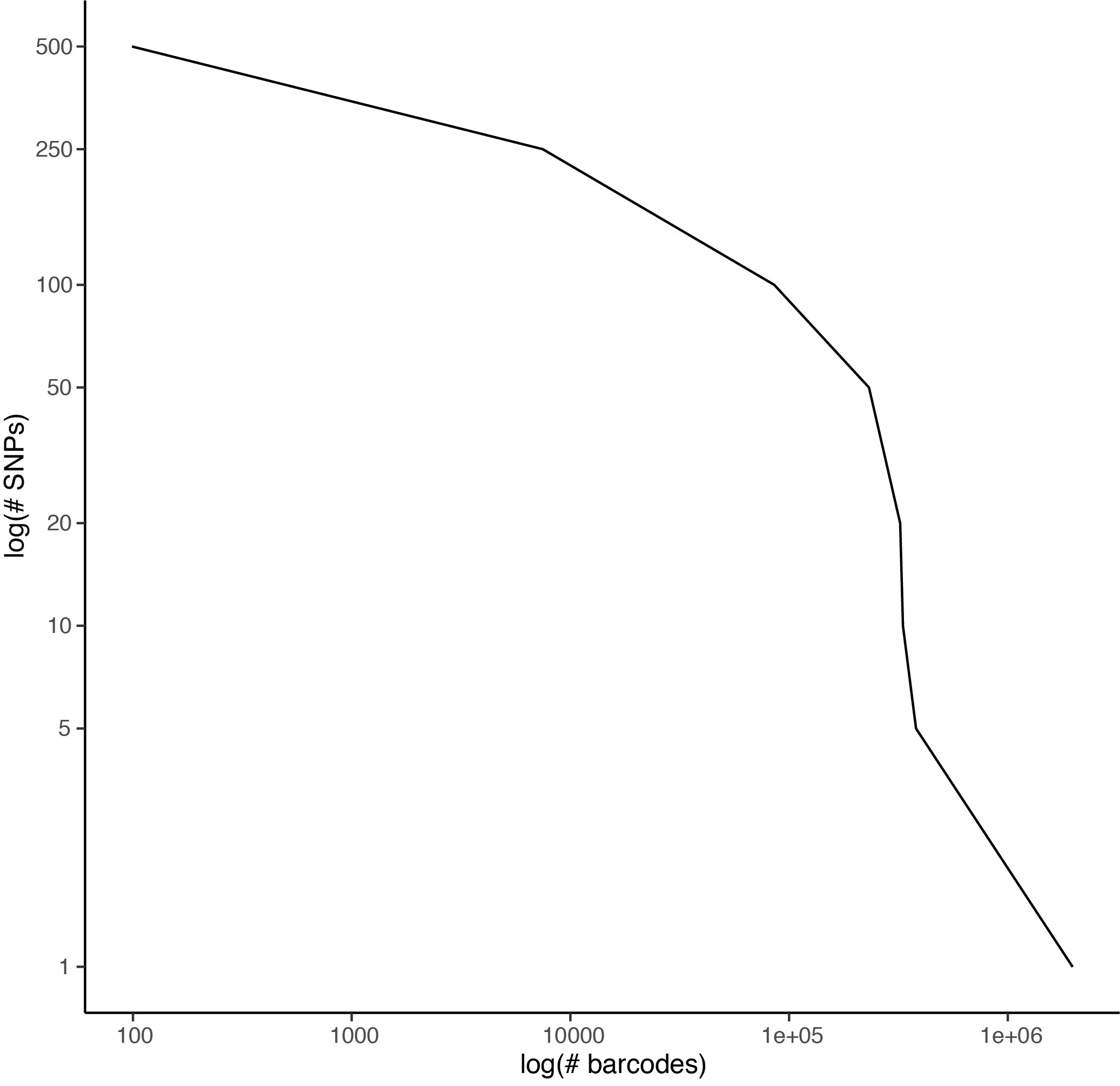

**Figure S1. Number of cell-containing barcodes per number of SNPs.**

We estimated the number of cell-containing droplets (x-axis) by the number of SNPs used by demuxlet to identify the donor of origin (y-axis).

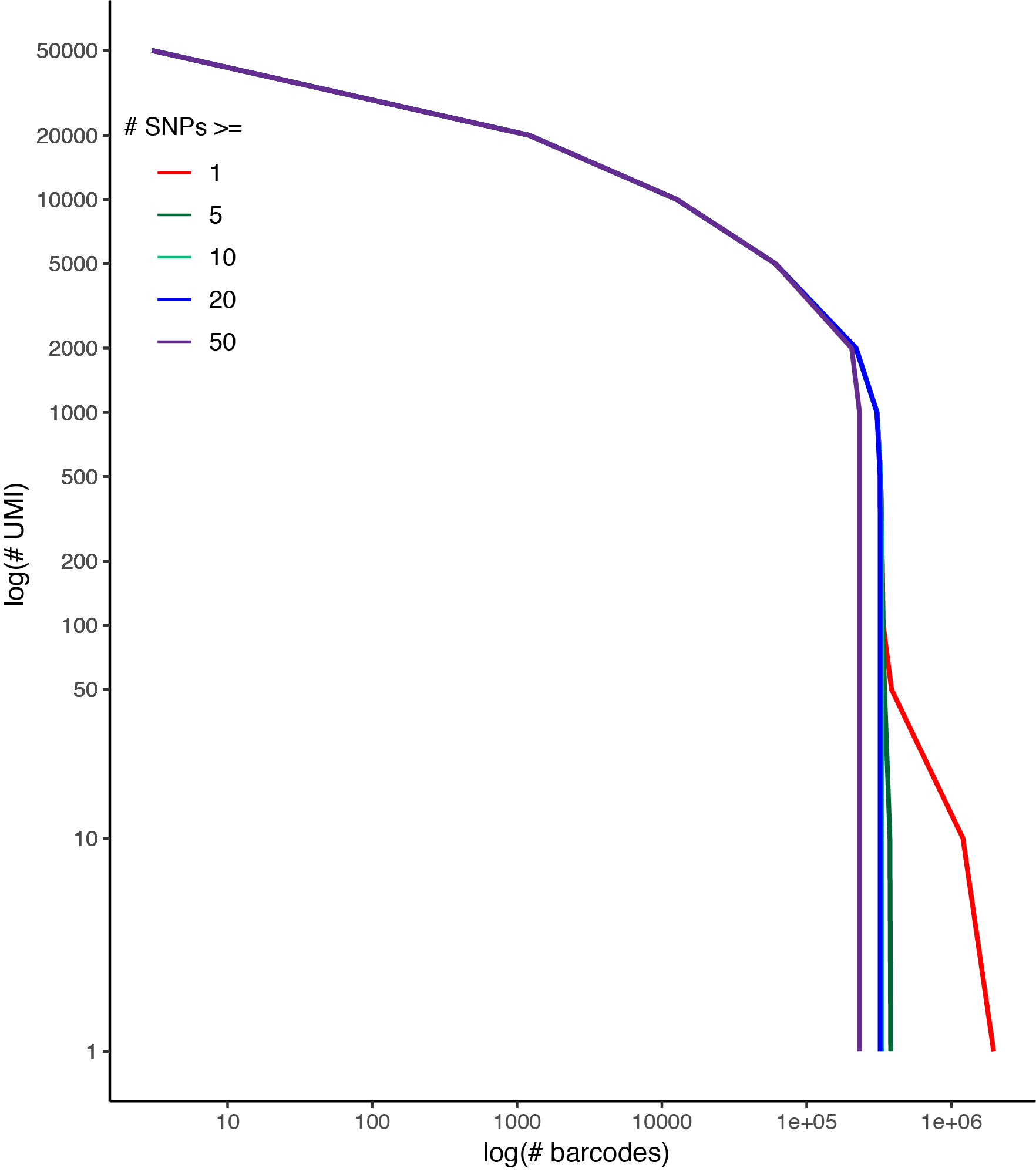

**Figure S2. Number of cell-containing barcodes per number of UMIs.**

We estimated the number of cell-containing droplets (x-axis) by the number of UMIs filtered for one (red), five (green), 10 (blue), and 50 (purple) SNPs used by demuxlet to identify the donor of origin (y-axis).

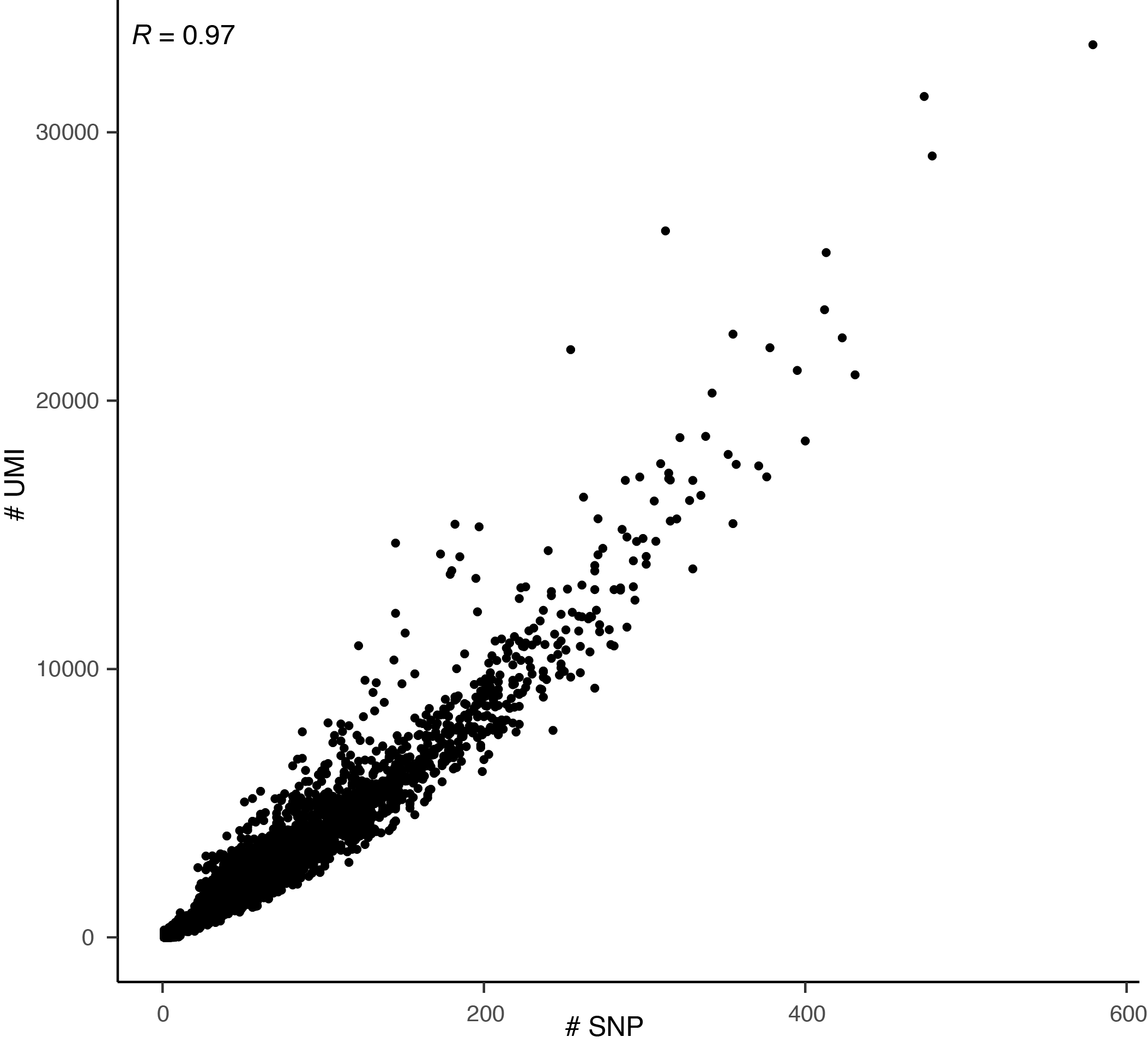

**Figure S3. Correlation of number of SNP and nUMIs.**

The number of SNPs (x-axis) and the number of UMIs (y-axis) estimated from demuxlet (Pearson *R*=0.97). Each point represented a cell-containing droplets, sampled to 20,000 cell-containing droplets.

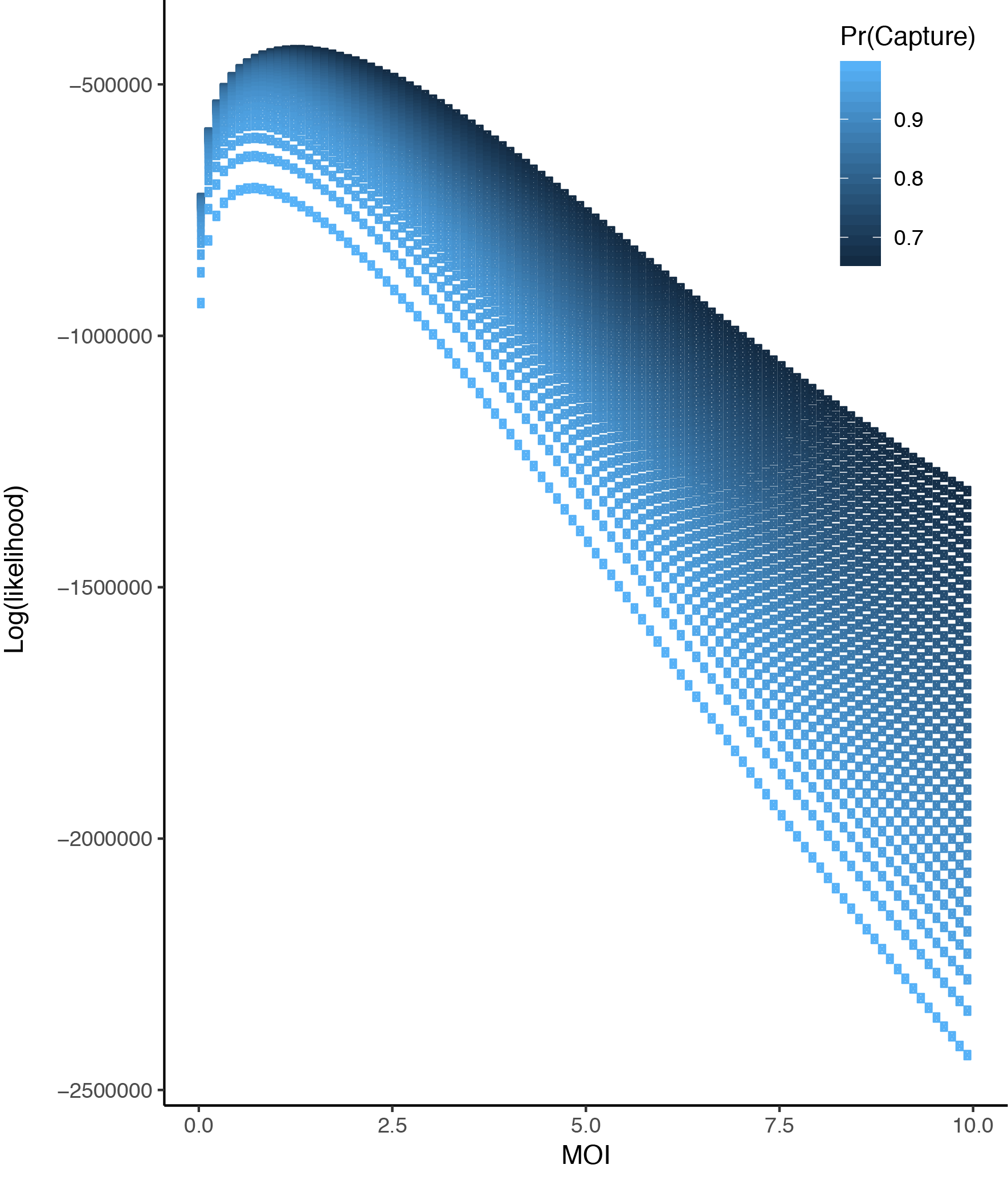

**Figure S4. MOI probability.**

Given a capture rate, we estimated the likelihood (y-axis) of seeing the transcript at a given MOI (x-axis).

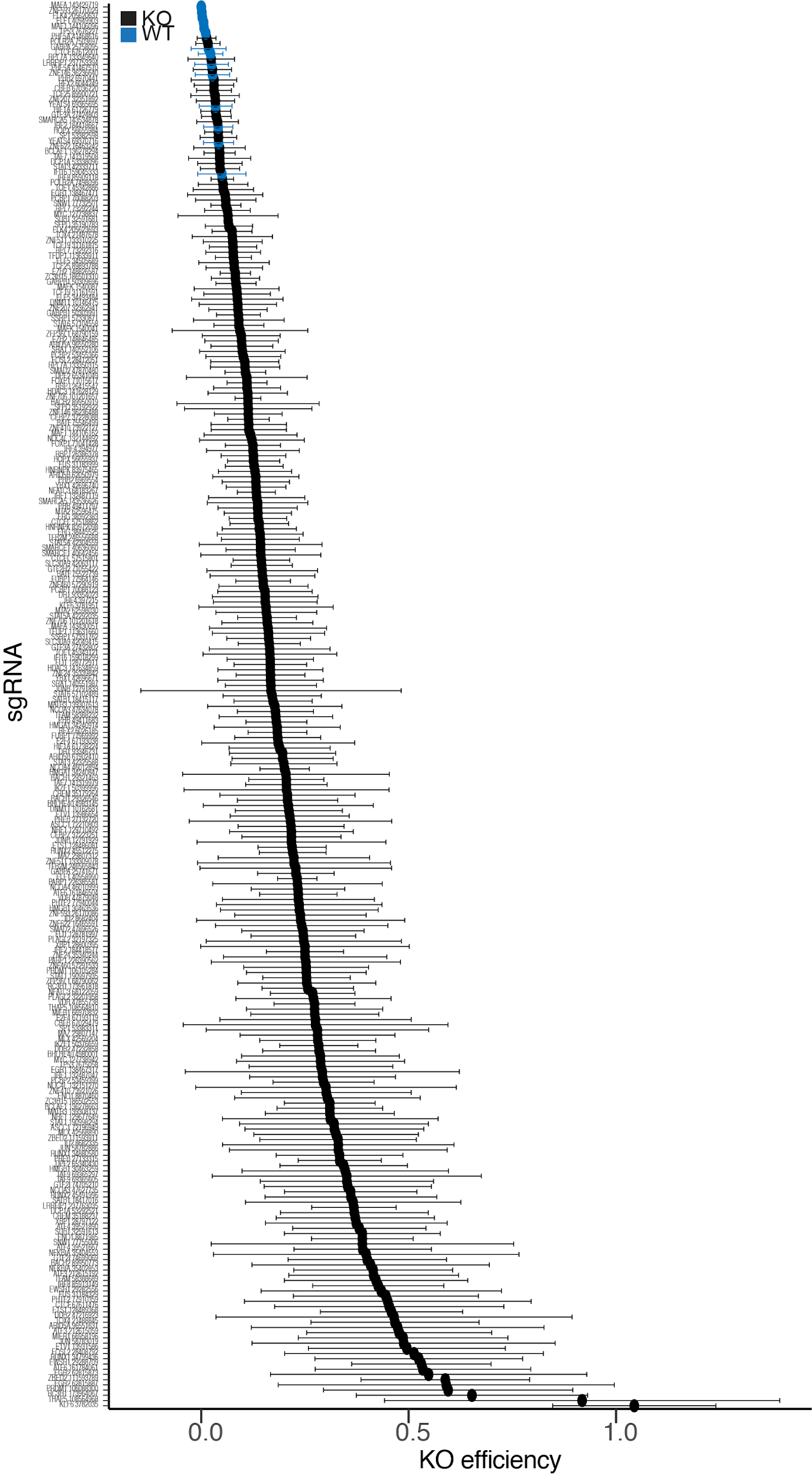

**Figure S5. sgRNA KO efficiency**

We estimated the average sgRNA KO efficiency (x-axis) per sgRNA (y-axis). Each point represents the average KO efficiency and error bars are the standard deviations across donors.

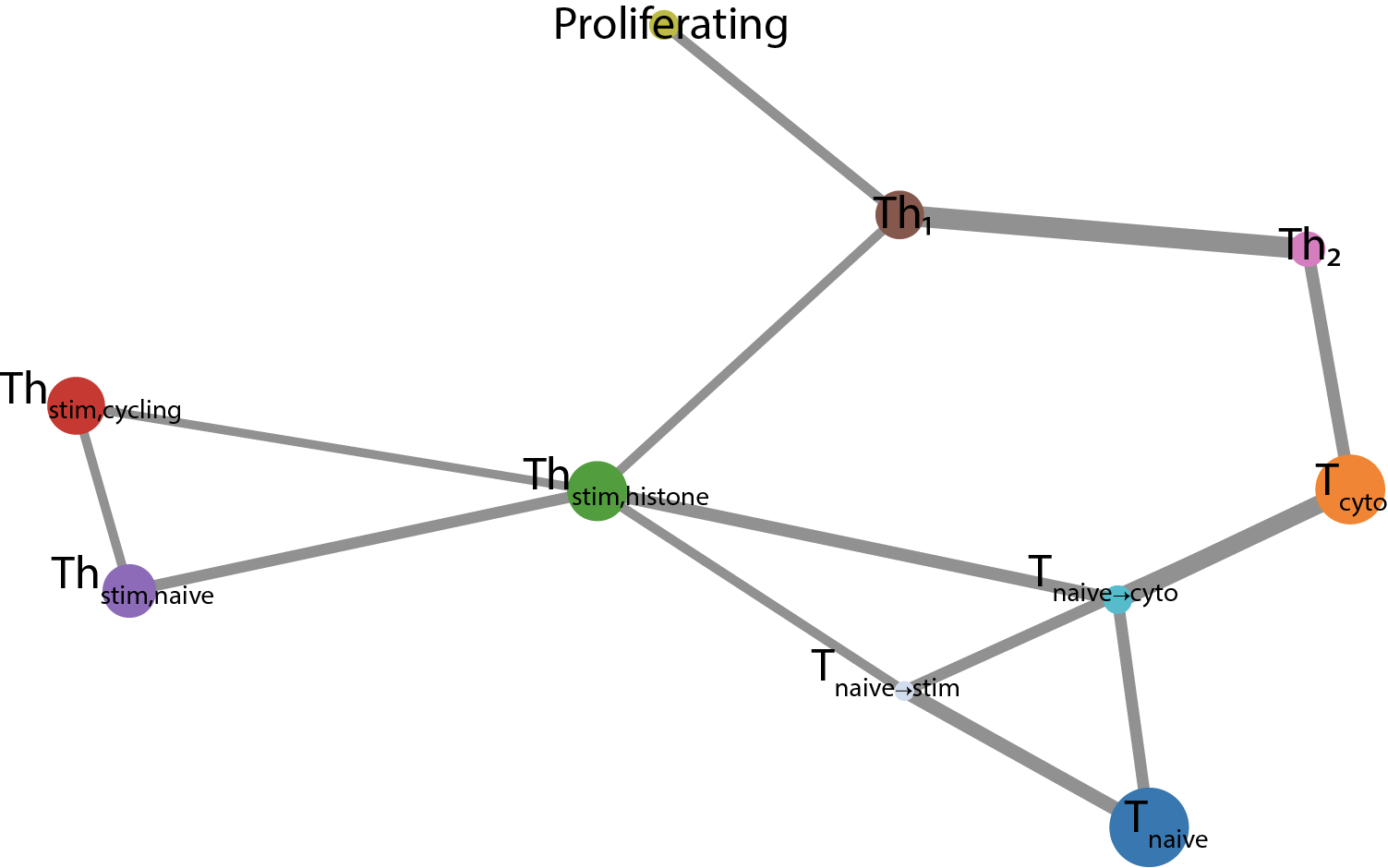

**Figure S6. Cell state trajectory.**

Cell state lineage trajectory using PAGA (*62*), where every node is a cell population and the size of the node corresponds to the size of the population. The width of each edge is the strength of connection between the nodes. The color of the point corresponds to the population in Fig. 2A.

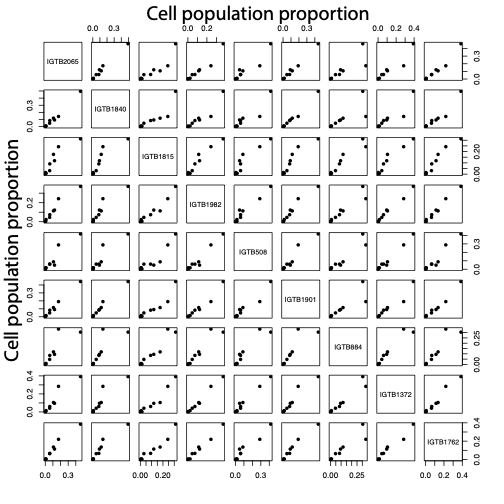

**Figure S7. Cell proportion across donors.**

Correlation matrix (Pearson *R*) of cell type proportions across donors. Each point represents a cell type.

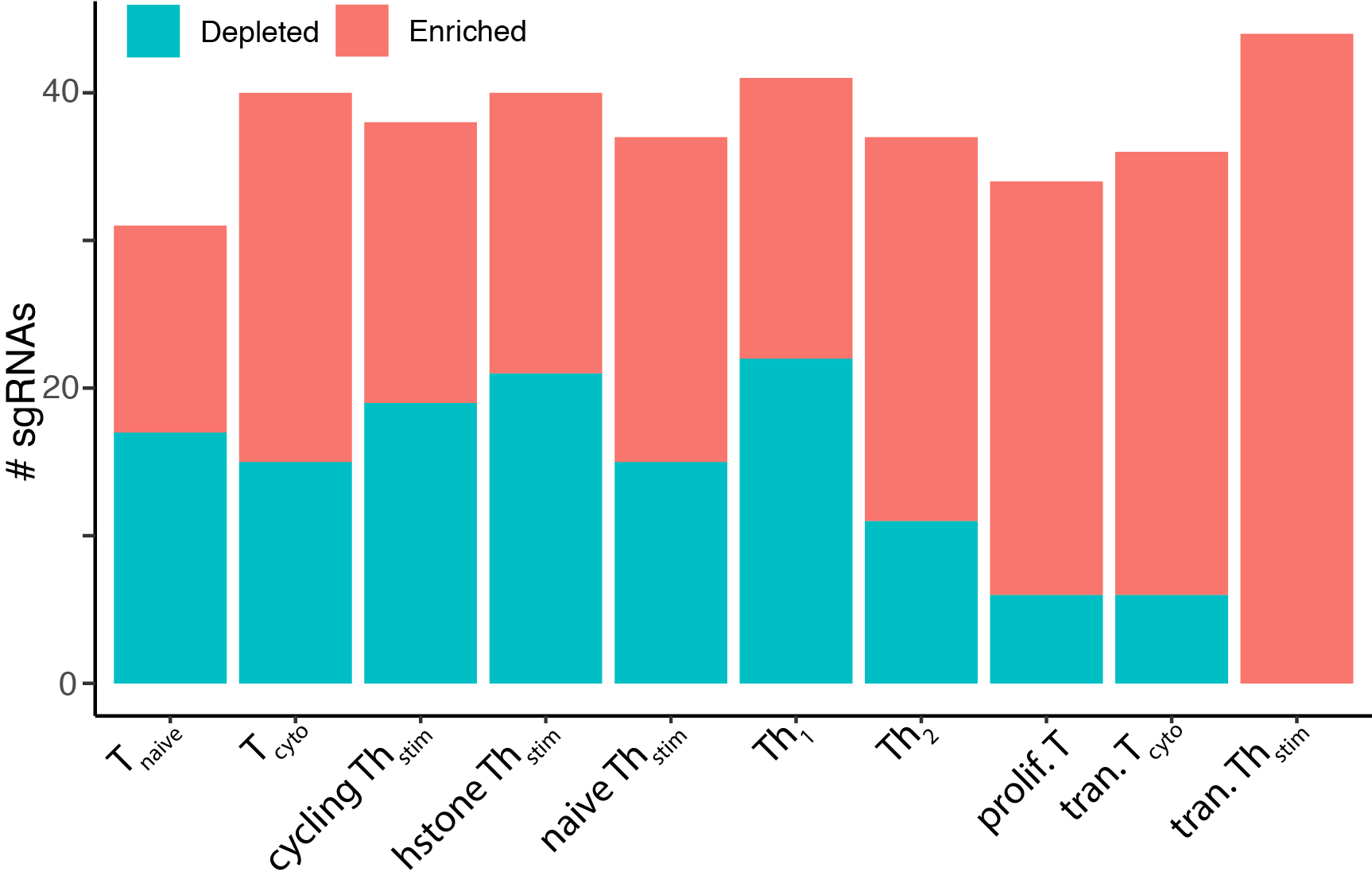

**Figure S8. sgRNA enrichment and depletion in a cluster.**

For each cluster, we calculated if the proportion of cells belonging to sgRNA is enriched (z-score > 1.5) or depleted (z-score < -1.5) in each cluster as compared to the proportion of all cells belonging to that cell state.

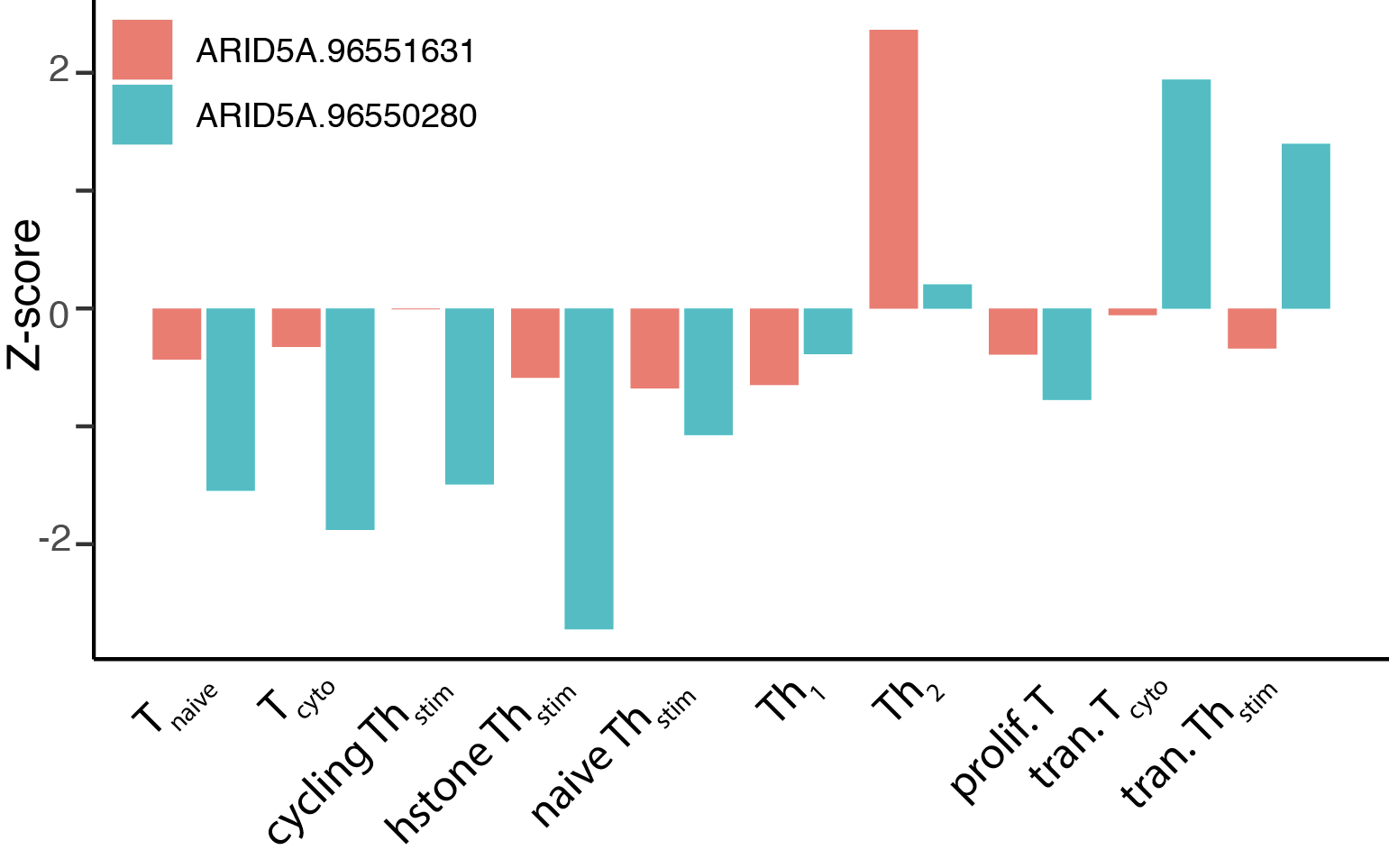

**Figure S9. ARID5A cell state enrichment and depletion**

For each cluster, we calculated if the proportion of cells belonging to both *ARID5A-targeting* sgRNAs (ARID5A, cutsite: chr2:96551631 in pink and chr2:96550280 in blue) and calculated a z-score (y-axis) for each cell state (x-axis).

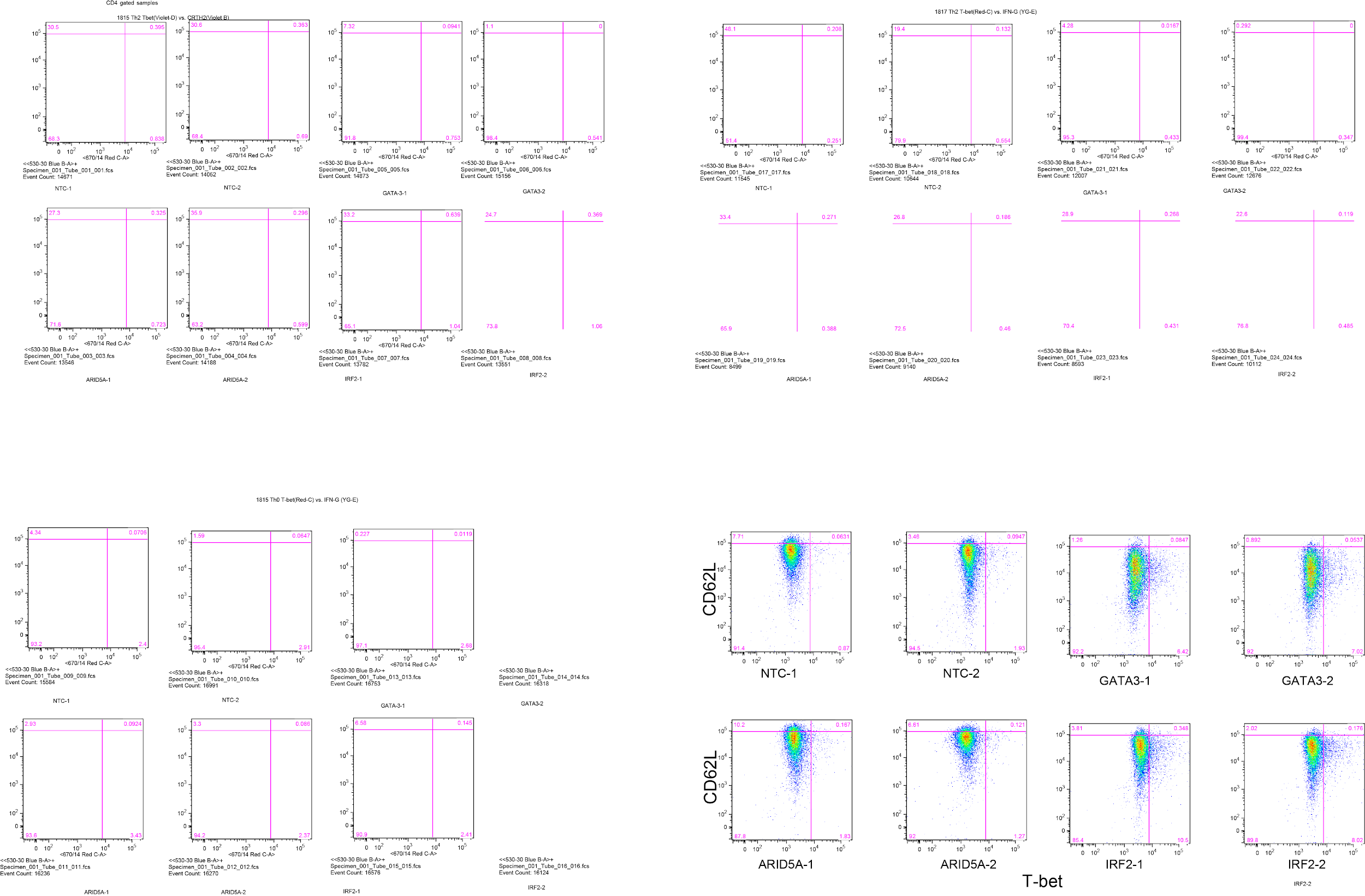

**Figure S10. Th_1_ *vs.* Th_2_ validation**

Using FACS, we sorted for CD62L^+^ (Th_2_ marker, y-axis) and T-bet^+^ (Th_1_ marker, x-axis) cells in our activated CD4^+^ stimulation across 2 non-targeting controls (top left), *GATA3-targeting* (top right), *ARID5A-targeting* (bottom left), *IRF2-targeting* (bottom right) sgRNAs.

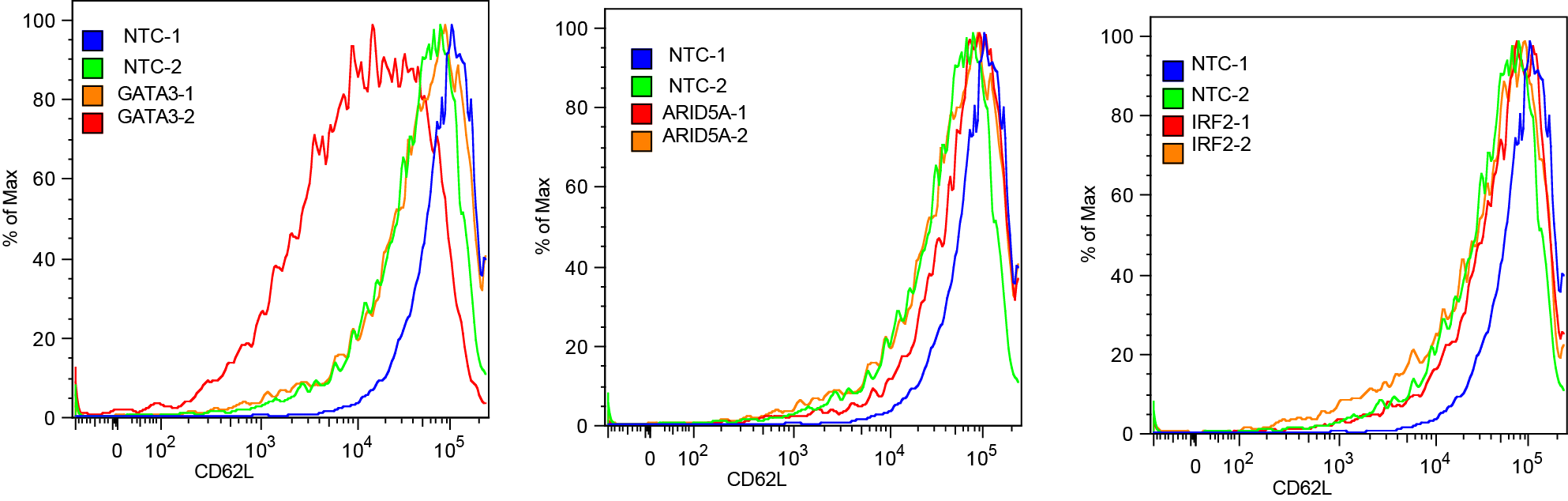

**Figure S11. Th_2_ polarization**

Under Th_2_ stimulation condition, we sorted for proportion of CD62L^+^ (Th_2_ marker, x-axis) across our eight sgRNA conditions. Green and blue are non-targeting controls compared to our red and orange knockout sgRNAs (*GATA3-targeting* sgRNAs left panel, *ARID5A-targeting* sgRNAs in middle panel, *IRF2-targeting* sgRNAs right panel)

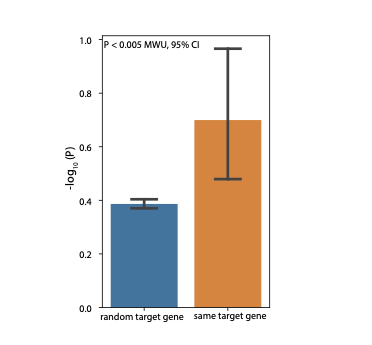

**Figure S12. Number of TF interaction downstream genes per TF.**

Difference between average -log_10_(*P*) for agreement of downstream genes for randomly paired sgRNAs (right) and sgRNAs targeting the same regulator. Agreement of downstream genes for a pair of sgRNAs was tested using a Chi-squared test, and the difference in the -log_10_(*P*) distribution was tested using Mann-Whitney U-test.

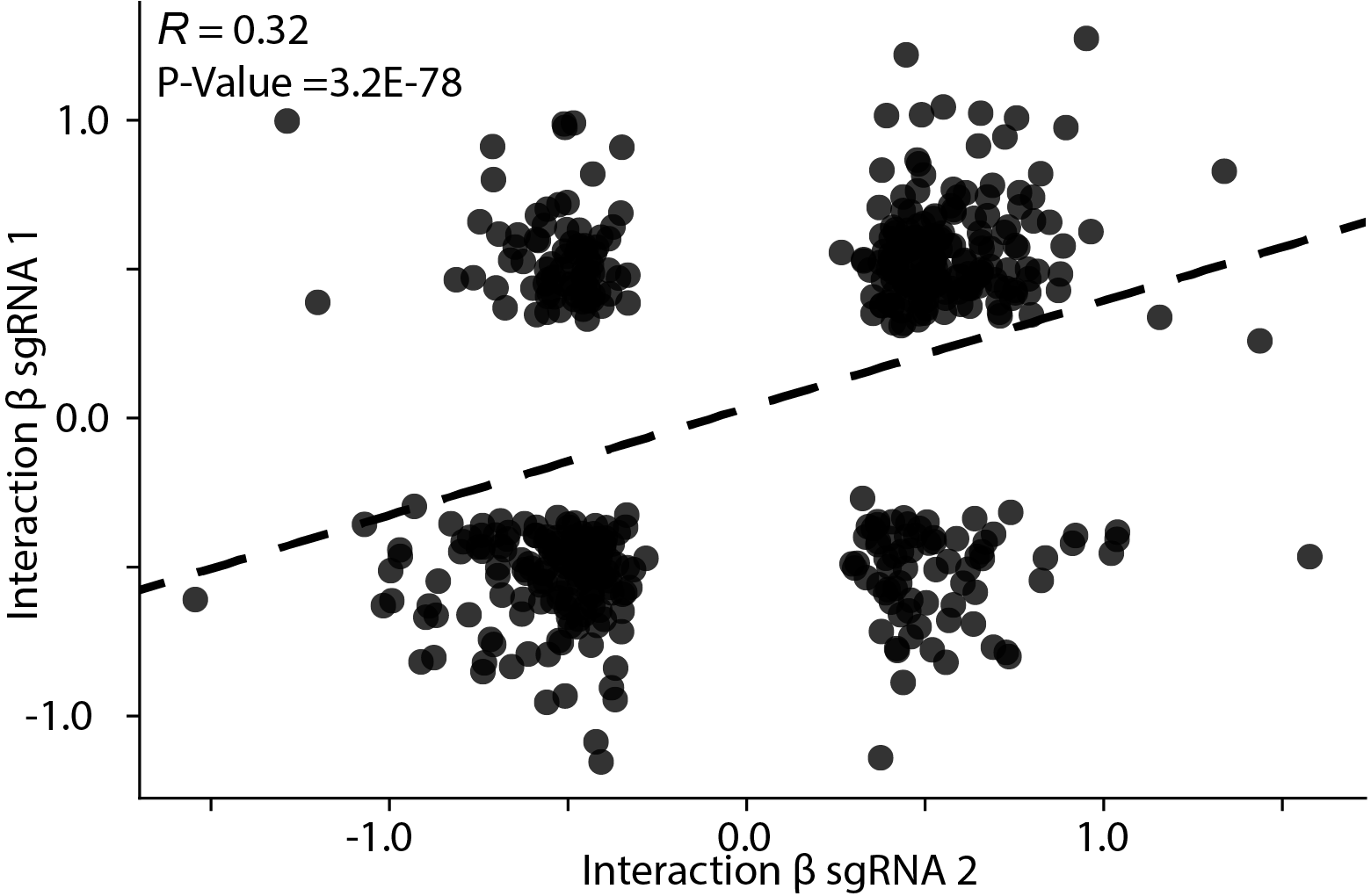

**Figure S13. Interaction effect size correlation.**

The correlation of interaction effect sizes for both sgRNAs targeting the same gene, sampled to 500 points. Each point is a sgRNA pair, with sgRNA 1 on the y-axis aond sgRNA 2 on the x-axis. The dashed line is the trend line.

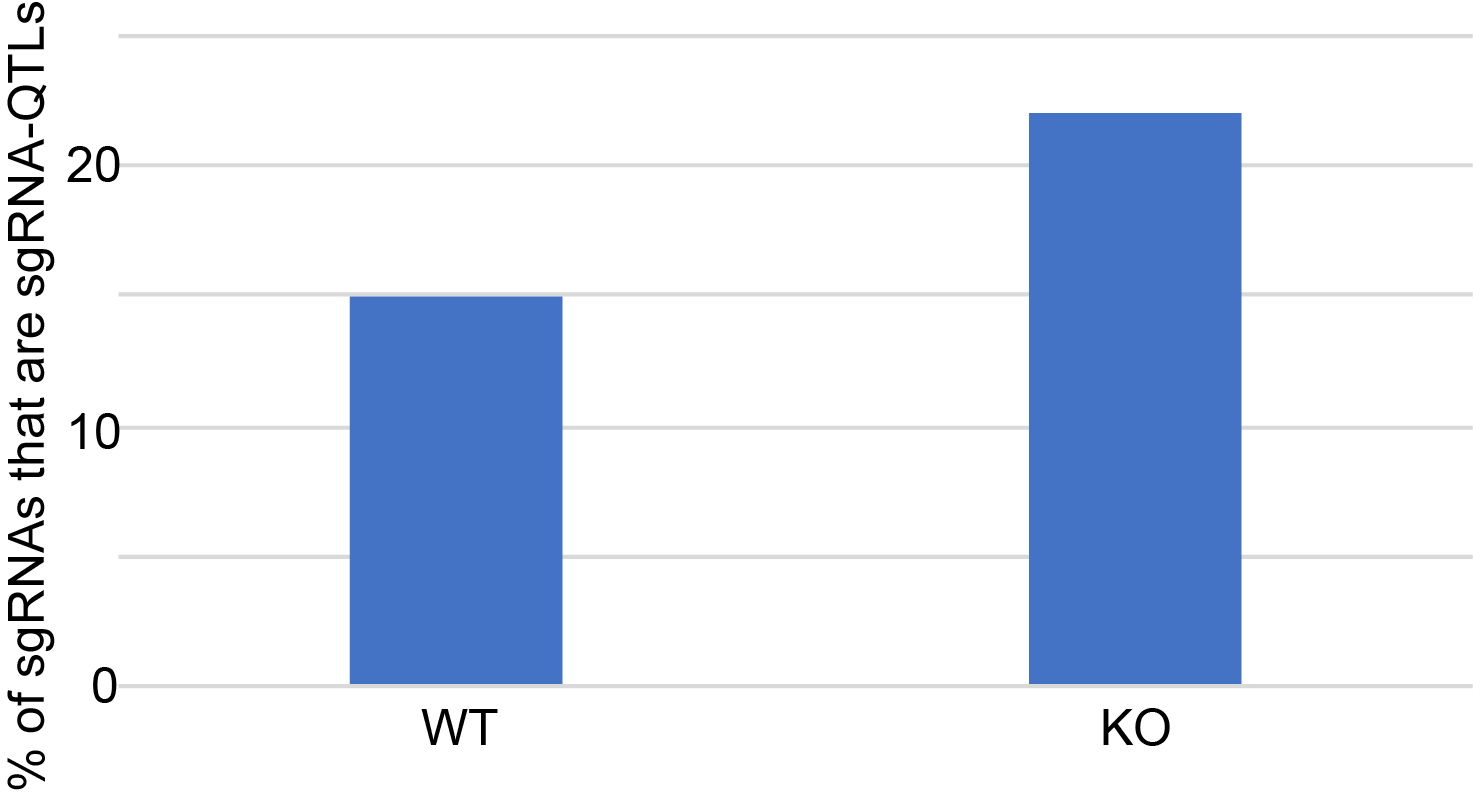

**Figure S14. Percentage of of sgRNAs that have an eQTL.**

Percentage (y-axis) of total WT (14) and KO (244) sgRNAs (x-axis) that have an eQTL.

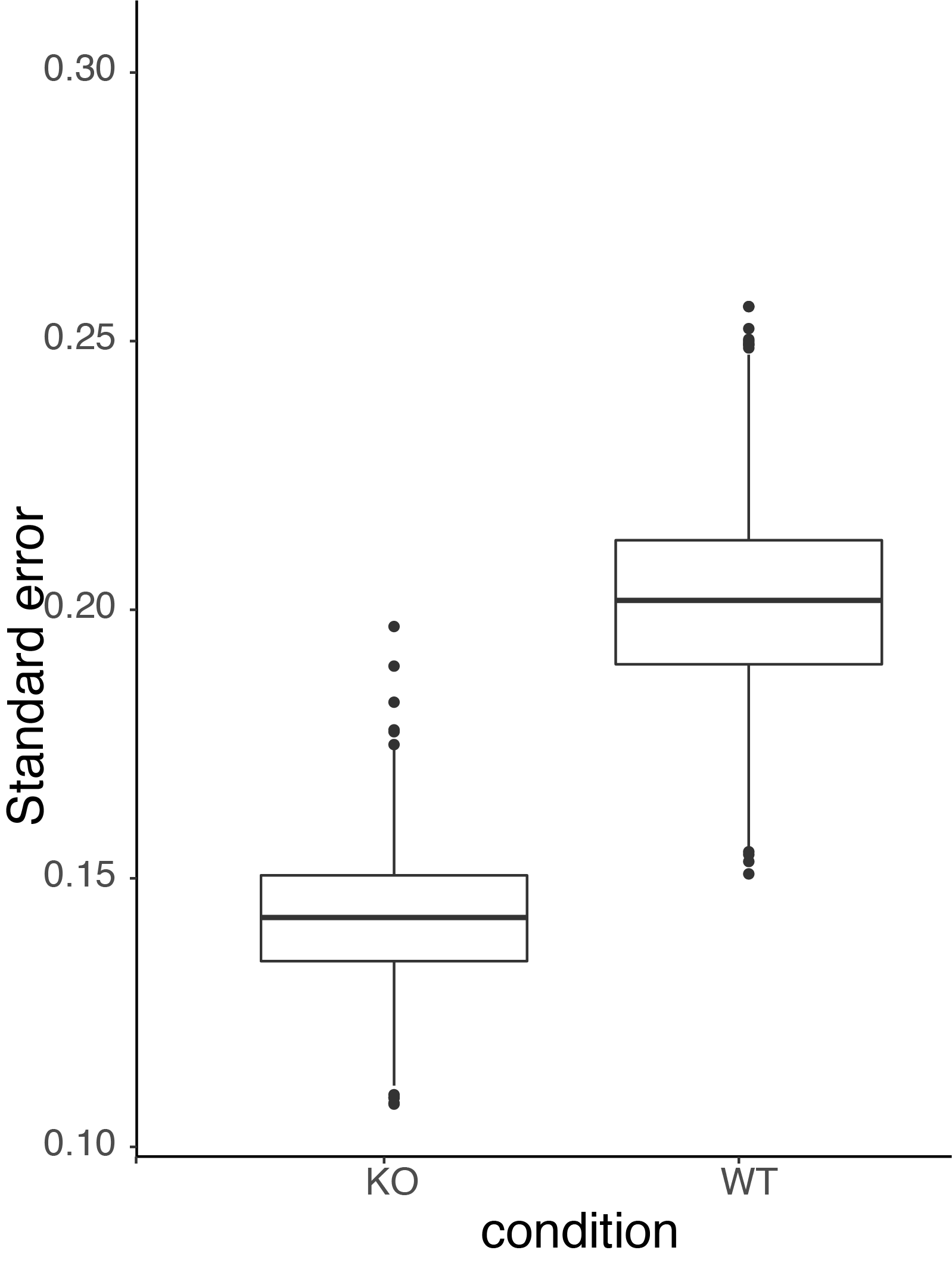

**Figure S15. Standard error simulation.**

The standard errors on 1,000 simulations with an effect size = 0.5, including a regulator variable (WT) and excluding the regulator variable (KO).

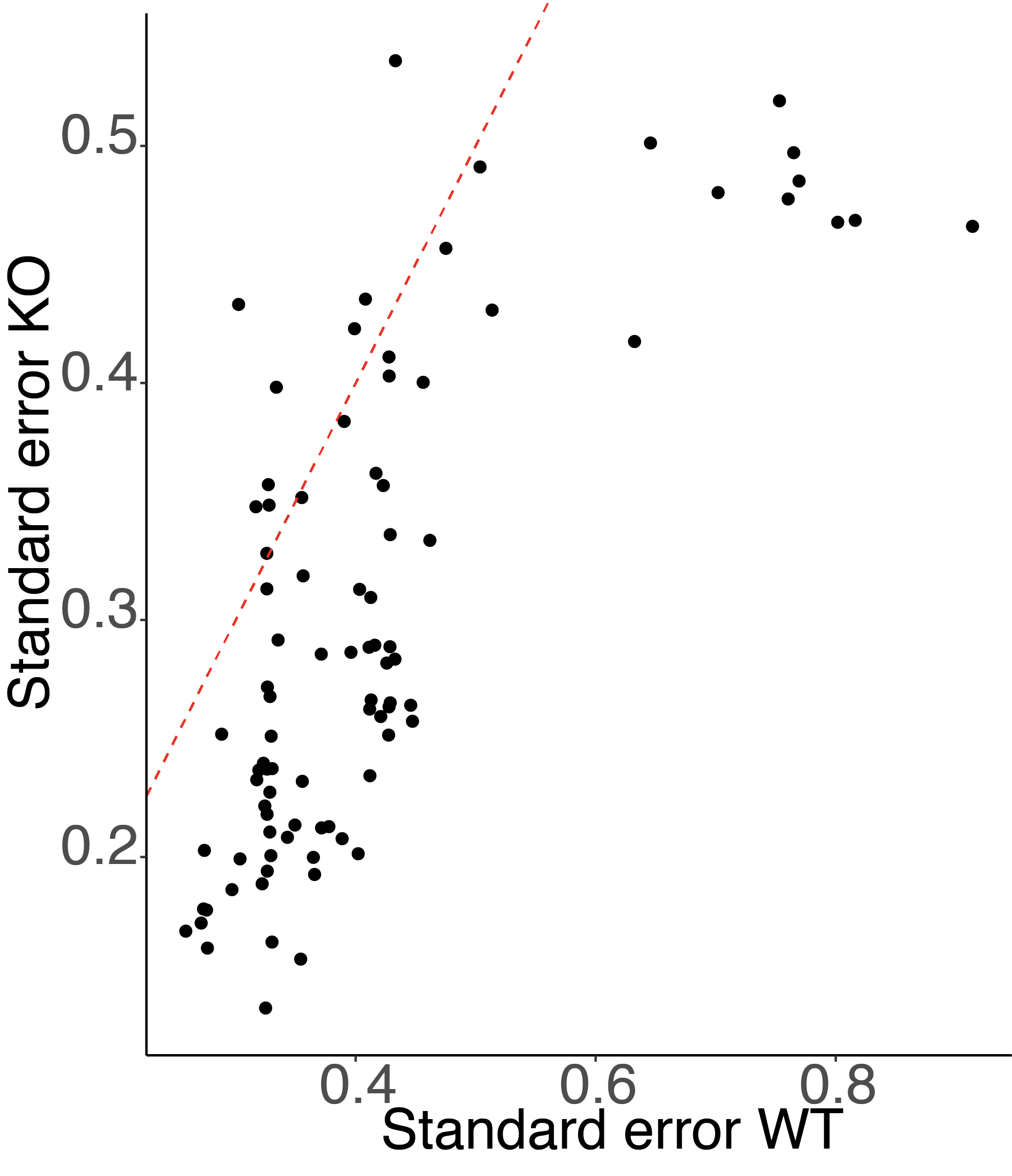

**Figure S16. Standard error of eQTLs.**

Standard error of eQTL (y-axis) compared to the standard error of the association in a WT sgRNA (x-axis). The red dashed line is an abline(slope=1, intercept=0)

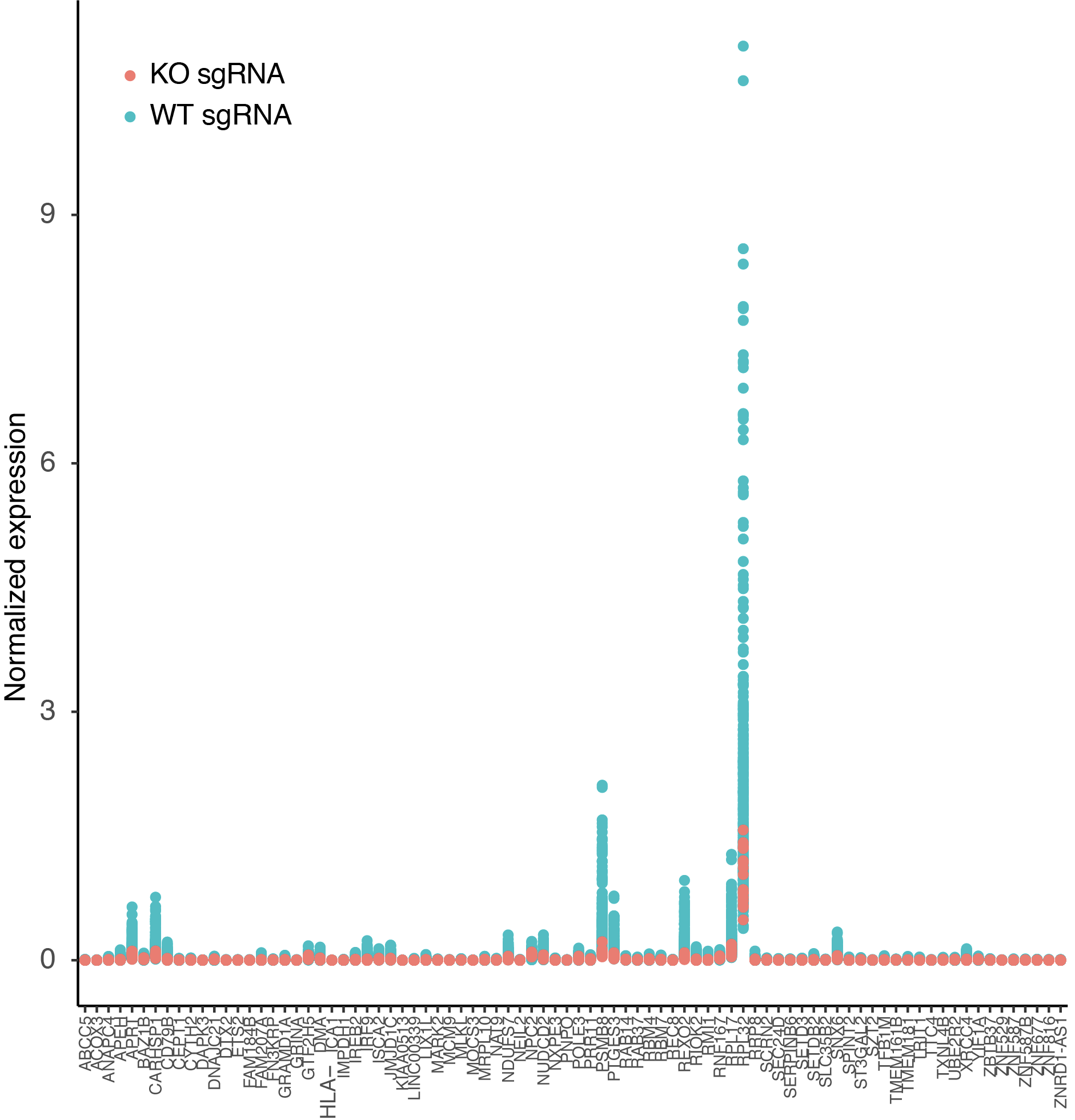

**Figure S17. eGene expression.**

Normalized expression (y-axis) of eGenes (x-axis) in the KO sgRNA (pink) and 14 WT sgRNAs (blue).

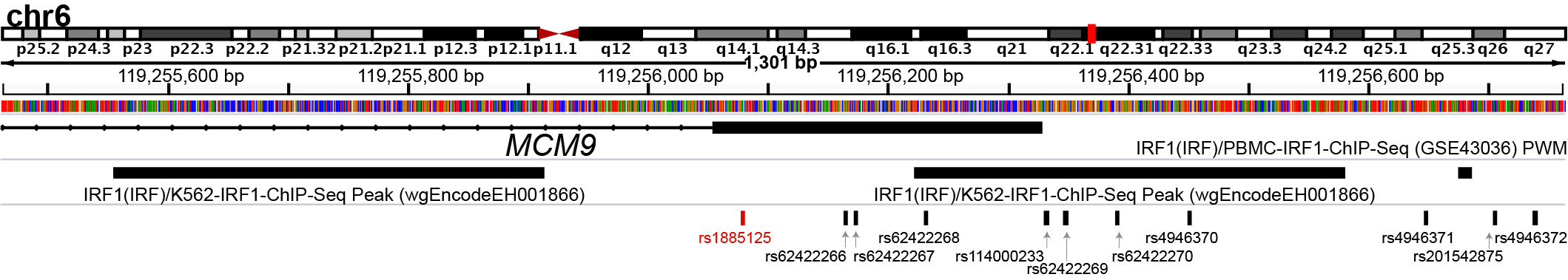

**Figure S18. Track of *MCM9* locus.**

Shown is a 1.3 kb window around the first *MCM9* exon, with three epigenetic annotations surrounding the region. There are two IRF1 ChIP-Seq peaks in K562 (IRF1(IRF1)/K562-ChIP-Seq Peak (wgEncodeEH001866)) and one from IRF1 position weight matrix calculated from a peripheral blood mononuclear cell ChIP-Seq (IRF1(IRF1)/PBMC-IRF1-ChIP-Seq (GSE43036) PWM). Below are SNPs in LD (D’ >= 0.97) with our eQTL (in red).
